# Microtubule organization and molecular architecture of ciliary basal bodies in multiciliated airway cells

**DOI:** 10.1101/2025.09.04.674302

**Authors:** Emma J. van Grinsven, Eugene A. Katrukha, Karen B. van den Anker, Fangrui Chen, Jeffrey M. Beekman, Lukas C. Kapitein, Anna Akhmanova

## Abstract

Microtubule organization depends on cell type and function. Microtubule networks of many differentiated cell types, such as epithelial cells, are poorly understood due to their high density. Here, we used expansion microscopy to quantitatively describe the three-dimensional organization of the microtubule network in human airway multiciliated cells. In these cells, most apical and apicobasal microtubules nucleate and anchor at the basal foot, a part of the ciliary basal body. Using a newly developed volumetric averaging tool, we generated a high-resolution 3D map of the basal body, delineating the position of structural components and proteins involved in microtubule nucleation and anchoring. γ-TuRC, its binding partners NEDD1 and Augmin/HAUS, and centriolar appendage proteins ninein and AKNA localize to the basal foot. Functional analyses demonstrated that NEDD1, but not ninein or HAUS are essential for basal foot-dependent microtubule organization. Our data reveal the distinct architecture of microtubule-organizing centers responsible for the formation of dense microtubule arrays in multiciliated cells.

## Introduction

Microtubules play an essential role in determining cell morphology and function. Therefore, investigating how microtubule networks are established and maintained in various cell types is important for understanding cell differentiation (reviewed in (Akhmanova and Kapitein, 2022; Muroyama and Lechler, 2017; Sanchez and Feldman, 2017; van Grinsven and Akhmanova, 2025). In columnar epithelial cells, microtubules typically extend from the apical to the basal side to control asymmetric intracellular transport (Bacallao et al., 1989; Kunimoto et al., 2012; Mogensen et al., 1989; Yano et al., 2013). Such microtubule organization is also present in multiciliated epithelial cells (MCCs) present in airways, reproductive ducts and brain ventricles, where hundreds of cilia project from the centriole-like structures called basal bodies and beat in a coordinated manner to create fluid flow (Clare et al., 2014; Lyu et al., 2024; Tateishi et al., 2017). Coordinated ciliary beating in such epithelia depends on planar cell polarity and the organization of the apical cytoskeleton, which in MCCs consists of a complex network of actin filaments, intermediate filaments and microtubules (Clare *et al*., 2014; Gordon, 1982; Herawati et al., 2016; Kunimoto *et al*., 2012; Nakayama et al., 2021; Shiratsuchi et al., 2024; Takagishi et al., 2020; Tateishi *et al*., 2017; Usami et al., 2021; Vladar et al., 2012; Werner et al., 2011). Dysfunction of MCCs leads to human diseases such as primary ciliary dyskinesia (Fliegauf et al., 2007).

Although the importance of microtubules for cellular function is well-established, we currently miss a comprehensive three-dimensional view of the dense polarized microtubule networks in differentiated epithelial cells. Conventional immunofluorescence-based analyses provide important insights but are insufficient to resolve the network architecture (Goldspink et al., 2017; Toya et al., 2016). Electron microscopy (EM) represents a powerful alternative, demonstrated by studies of whole mitotic spindles (Kiewisz et al., 2022; Redemann et al., 2017) and pancreatic beta cells (Muller et al., 2021). However, EM-based analyses are extremely laborious, difficult to combine with staining of specific proteins and are often confined to subcellular regions (e.g., (Clare *et al*., 2014)). Expansion microscopy overcomes these challenges (Chen et al., 2015; Wassie et al., 2019). It does not only permit imaging three-dimensional organization of cytoskeletal filaments and organelles but also allows to determine the molecular architecture of cellular structures that are below the resolution limit of conventional optical microscopy. For example, recent advances in expansion microscopy together with other super-resolution microscopy techniques provided a comprehensive map of centriolar proteins (Chang et al., 2023; Chong et al., 2020; Laporte et al., 2024; Woglar et al., 2022; Yang et al., 2018).

Super-resolution microscopy revealed important details of the structure of ciliary basal bodies and their appendages, basal feet, which are needed for ciliary positioning and synchronized beating (Liu et al., 2020; Lyu *et al*., 2024; Nguyen et al., 2020). Mutations in the gene encoding ODF2/cenexin, a structural component of centriolar appendages, abolished basal foot formation and resulted in the loss of apical microtubules (Ishikawa et al., 2005; Kunimoto *et al*., 2012; Tateishi et al., 2013). Other basal foot components include CEP112, centriolin, galectin-3, ninein, CEP170, NEDD1 and γ-TuRC (Clare *et al*., 2014; Nguyen *et al*., 2020). EM-based analysis of the apical section of MCCs showed that microtubules associate with the basal foot, and this was impaired in a mouse knockout of galectin-3 (Clare *et al*., 2014). However, a detailed understanding of the proteins required to establish and maintain an apicobasal microtubule network in MCCs is lacking.

Here, we used expansion microscopy combined with computational techniques to map the complete microtubule network as well as the ultrastructure of the microtubule organizing centers (MTOCs) of airway MCCs. First, we optimized the workflow of Ten-fold Robust Expansion Microscopy (TREx)(Damstra et al., 2022) to reliably expand human airway epithelium. We quantitatively analyzed the microtubule network and microtubule subsets in MCCs during maturation and confirmed that most microtubules from the basal foot. By automatically averaging hundreds of basal bodies, we generated an ultrastructural map of the basal body, analyzed its similarities and differences with centrioles and identified candidate proteins responsible for MTOC activity at the tip of the basal foot. Lastly, we investigated the function of these proteins and found that NEDD1 but not HAUS or ninein are essential for γ-TuRC localization at the basal foot and apical microtubule anchoring. Together, this work provides insight into the architecture of elaborate microtubule arrays in MCCs and the distinct composition of their MTOCs.

## Results

### Characterization of microtubule networks in airway epithelium by expansion microscopy

To study human nasal airway epithelial cells (HNECs), we differentiated stem cells, called basal cells, using an air-liquid interface (ALI) culture system (Fig. 1A,B)(Gray et al., 1996; Rodenburg et al., 2023). Within 15 days at ALI, we observed a tightly packed, pseudostratified cell layer containing three major cell types: proliferative cells at the basal side, secretory cells with MUC5AC-containing granules, and MCCs marked by β-IV tubulin-positive cilia (Fig. 1B,C)(Deprez et al., 2020). MCCs were apparent already at the onset of differentiation, at ALI culture day 5, and their number increased and stabilized over time (Fig. 1D). At the onset of differentiation, MCCs generate large numbers of basal bodies through deuterosome-dependent amplification; basal bodies then dissociate from the deuterosome, migrate towards the apical surface and subsequently organize into floret, random and aligned patterns (Al Jord et al., 2014; Herawati *et al*., 2016; Klos Dehring et al., 2013; Shiratsuchi *et al*., 2024). Using stimulated emission depletion (STED) microscopy of basal body and basal foot markers ODF2 and centriolin, respectively, we could distinguish all these patterns and observe a gradual transition from amplification to alignment of basal bodies (Fig. 1E,F).

**Figure 1:**
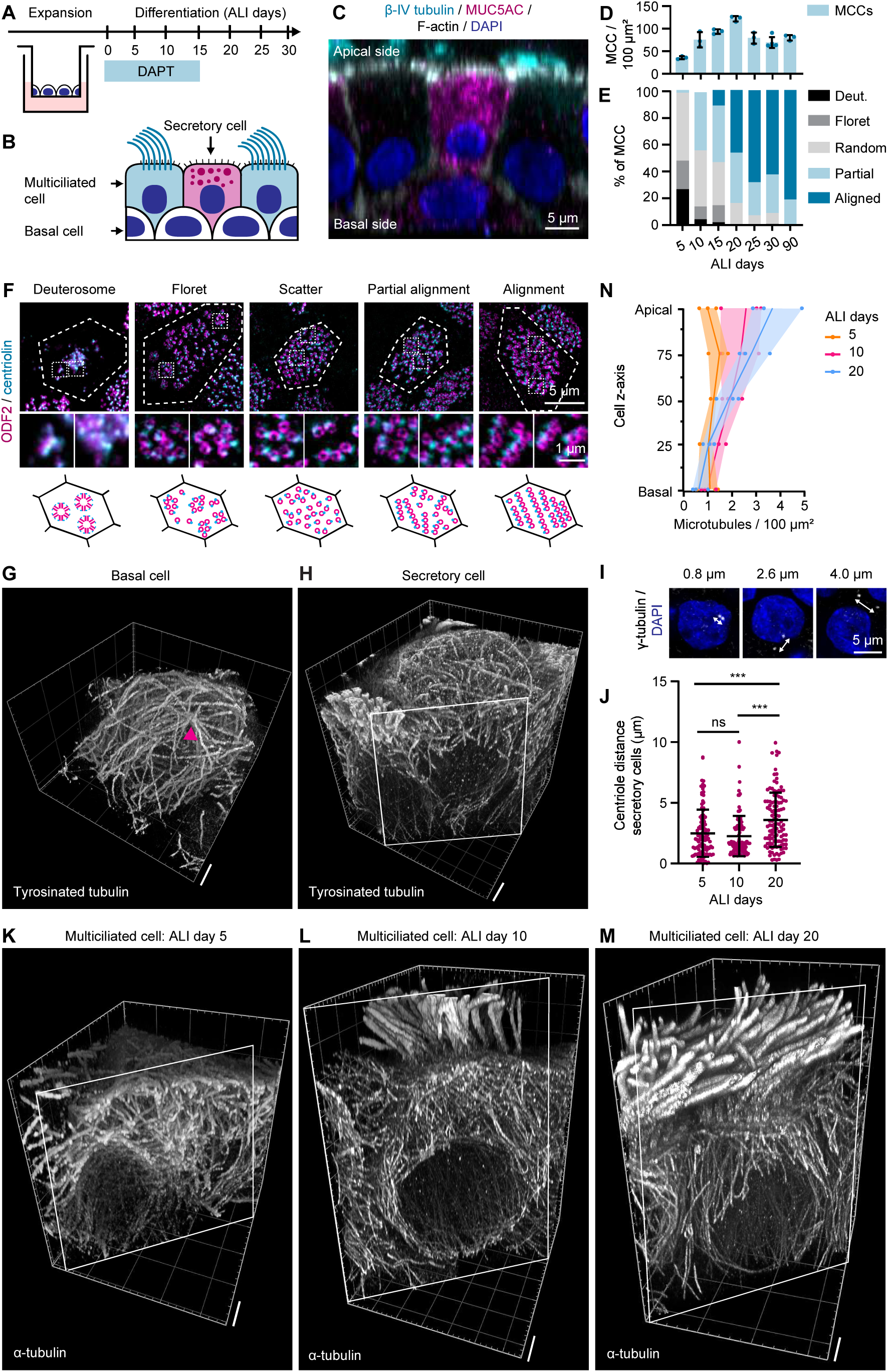
Microtubule network of airway epithelial cells imaged using TREx. (A) Scheme of air-liquid interface (ALI) culture of HNEC. Basal cells are seeded and expanded on ALI until confluent; cells are differentiated for 15 days in the presence of Notch inhibitor DAPT and subsequently cultured on ALI. (B) Scheme of pseudostratified airway epithelium with multiciliated cells (MCCs) (blue), secretory cells (magenta) and basal cells (white). (C) Side-view of pseudostratified HNECs stained for β-IV tubulin (cyan), MUC5AC (magenta), F-actin (white) and DAPI (blue). (D) Number (mean±SD) of MCCs per 100μm² at different days of differentiation on air-liquid interface (ALI). Three 125 μm² fields of view were analyzed per timepoint. (E) Quantification of MCCs during differentiation on air-liquid interface (ALI) categorized into deuterosome (Deut.), Floret, Random, Partial aligned (Partial) and Aligned basal body pattern. n, number of analyzed cells: 5 days, n=52; 10 days, n=64; 15 days, n=89; 20 days, n=78; 25 days, n=78, 30 days, n=74; 90 days, n=88. (F) MCCs with characteristic basal body patterns stained for ODF2 (magenta) and centriolin (cyan) acquired using STED microscopy. Dashed lines show cell outlines. Boxes mark regions enlarged in zoom. Bottom row represents schematics of characteristic basal body patterns. (G,H) Volumetric 3D renders of TREx images of microtubule network in a basal cell (G) and a secretory cell (H) stained for tyrosinated tubulin (white). Pink arrowhead highlights the MTOC. Scale bar (corrected for expansion) is ∼2μm. (I) Secretory cells stained for γ-tubulin (white) and DAPI (blue) in MCCs showing variation in centriole distance. (J) Distance (mean±SD) between two centrioles measured from center-to-center in secretory cell at ALI day 5, 10 and 20. Points represent one centriolar pair. n, number of analyzed cells: 5 days, n=108; 10 days, n=120; 20 days, n=115. (K, L, M) Volumetric 3D renders of TREx images of microtubule network in MCCs at ALI day 5 (K), ALI day 10 (L) and ALI day 20 (M) stained for α-tubulin (white). Scale bar (corrected for expansion) is ∼2μm. (N) Microtubule density map along the cell z-axis of MCCs at ALI day 5, 10 and 20. Points represent number of microtubules per cross-section of one cell. n, number of analyzed cells: ALI day 5, 10 and 20, n=3.

To explore microtubule networks of the three major airway cell types, we visualized them using TREx (Damstra *et al*., 2022). At ALI culture day 15, basal cells had a radial microtubule organization (Fig. 1G, MTOC indicated with an arrow, Movie 1). In contrast, in secretory cells, microtubules were distributed randomly (Fig. 1H, Fig. S1A, Movie 2), and although two γ-tubulin-positive centrosomes were present in each cell (Fig. 1I), they did not serve as focal points for microtubule organization (Fig. S1A). The distance between centrosomes in secretory cells increased in older cultures (Fig. 1I,J), possibly due to increased number of mucin granules (Rogers, 2003).

Microtubule organization in MCCs depended on their maturation (Fig. 1K-M). Basal bodies in MCCs were stained with a N-hydroxysuccinimidyl (NHS) ester dye, which labels primary amines of all cellular proteins, resulting in a signal proportional to the local protein density that provides contrast similar to electron microscopy (M’Saad and Bewersdorf, 2020). At ALI culture day 5, in MCCs undergoing basal body amplification and lacking cilia (Fig. S1B), microtubules organized radially around basal bodies (Fig. 1K, S1B, Movie 3). At ALI culture day 10, when in most MCCs cilia are present but appear short, microtubules started to align along the longitudinal cell axis (Fig. 1L, Movie 4). After ALI day 20, the apicobasal microtubule organization was established (Fig. 1M, Movie 5), consistent with earlier low-magnification two-dimensional observations of MCCs (Shiratsuchi *et al*., 2024; Tateishi *et al*., 2017; Tateishi *et al*., 2013).

We counted the number of microtubules at five MCC cross-sections, corrected for differences in cell area, and built a microtubule density map along the longitudinal axis (Fig. 1N, Fig. S1C,D). Whereas microtubules were distributed homogenously at ALI culture day 5, the overall microtubule density at the apical, but not at the basal side gradually increased during maturation. The share of vertical microtubules increased from ALI day 5 to ALI day 20, while the share of horizontal microtubules decreased (Fig. S1D,E). This rearrangement was especially prominent in the apical cross-section, where the share of vertically oriented microtubules increased from ∼6% to ∼67% (Fig. S1D,E). The number of basal bodies didn’t change (∼70-100 basal bodies per cell), and the average number of vertically oriented microtubules per basal body mildly decreased from 2.8 ± 0.6 at day 10 to 2.0 ± 0 at day 20 (n=3 cells). However, the apical area also decreased during maturation, and therefore, microtubule density increased (Fig. 1N).

In addition to the vertical, apicobasal network, MCCs form a horizontal meshwork just below the apical surface (Tateishi *et al*., 2017; Werner *et al*., 2011). Up to ALI culture day 10, apical microtubules were disorganized, but as the MCCs matured, they developed a regular microtubule mesh surrounding the basal bodies (Fig. 2A,B). After ALI culture day 20, MCCs contained microtubule bundles parallel to the apical cell membrane that were attached to basal bodies, supporting their proposed role in rotational polarity of basal bodies and planar cell polarity (Fig. 2C)(Nakayama *et al*., 2021; Werner *et al*., 2011). Microtubule 3D tracing using FIJI BigTrace plugin (Fig. S1F,G) showed that apical microtubule ends are clustered within the top 2% of cell height (Fig. 2G). Since we did not observe additional clusters of microtubule ends, we conclude that the majority of the microtubules are anchored at the apical side of the MCCs, in line with previous studies (Clare *et al*., 2014; Kunimoto *et al*., 2012; Nguyen *et al*., 2020; Tateishi *et al*., 2017).

**Figure 2:**
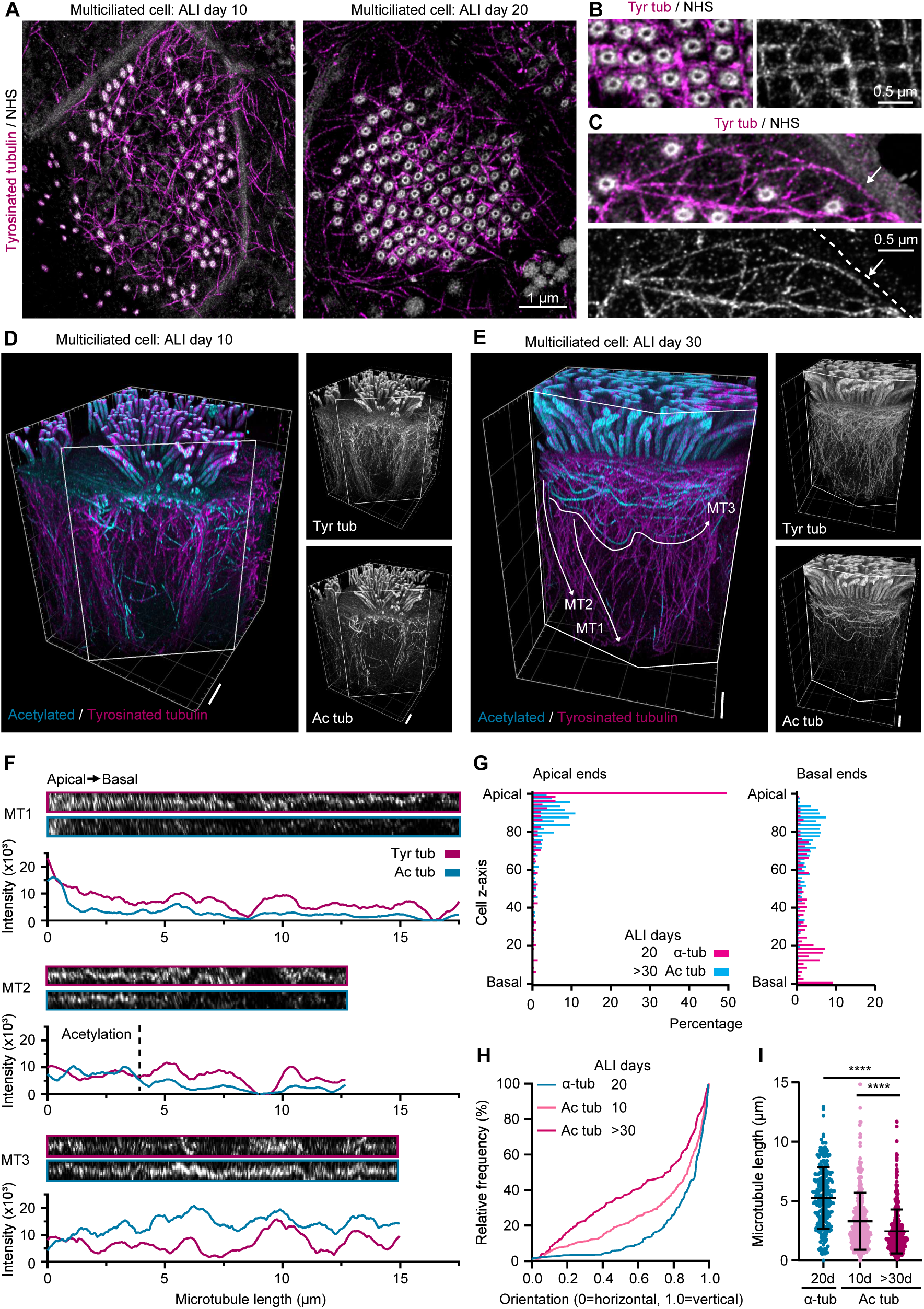
Visualization of post-translationally modified microtubule subsets in MCCs. (A-C) Apical microtubule meshwork of MCCs stained for tyrosinated tubulin (magenta) and NHS-ester dye (white) and imaged with TREx. Comparison between MCCs at ALI day 10 and 20 (A), bundles of microtubules in apical meshwork (B) and microtubules in close vicinity to a cell-cell junction (C). (D,E) Volumetric 3D renders of TREx images of microtubule network of MCCs at ALI day 10 (D) and day 30 (E) stained for acetylated (cyan) and tyrosinated (magenta) tubulin. Microtubule 3D traces in (E) were straightened and analyzed in (F). Scale bar (corrected for expansion) is ∼2μm. (F) Traced, straightened microtubules (highlighted in panel E) stained for acetylated (cyan) and tyrosinated (magenta) tubulin and their corresponding intensity profile (smoothened over 40 neighboring datapoints) from the apical (left) to basal (right) microtubule end. (G) Histogram of the position of apical (left) and basal (right) ends of microtubules analyzed using BigTrace plugin. Comparison between α-tubulin (cyan) of MCCs at ALI day 20 and acetylated tubulin (magenta) of MCCs at ALI day 30. n, number of traced cells: α-tubulin ALI d20, n=1, Ac tub ALI d30, n=3. (H) Cumulative frequency of microtubule orientation (0= horizontal, 1= vertical) in MCCs at ALI day 20 stained for α-tubulin, MCCs at ALI day 10 or 30 stained for acetylated tubulin. n, number of traced cells: α-tubulin ALI d20, n=1, Ac tub ALI d10, n=3, Ac tub ALI d30, n=3. (I) Length (mean±SD) of microtubule traces. Comparison between α-tubulin in MCCs at ALI day 20 and MCCs at ALI day 10 and 30 stained for acetylated tubulin. ****P<0.001 calculated using unpaired two-tailed Mann-Whitney U test. n, number of traced cells: α-tubulin ALI d20, n=1, Ac tub ALI d10, n=3, Ac tub ALI d30, n=3.

### Post-translational microtubule modifications in MCCs

Microtubules can be post-translationally modified over time (Janke and Magiera, 2020). Immunofluorescence staining showed that within cilia, tyrosinated and acetylated tubulin was present along the whole length, whereas detyrosination was only detected in long cilia and excluded from the ciliary tip (Fig. S2A-C). This detyrosination pattern reflects the absence of the B-tubule, which is more heavily detyrosinated than the A-tubule, from the distal end of the cilium (Chhatre et al., 2025; Johnson, 1998).

Within the cell body, we could readily detect acetylated, but not detyrosinated microtubules. In MCCs at ALI culture day 10, almost all acetylated microtubules were oriented vertically (Fig. 2D, Movie 6). In contrast, in mature MCCs (cultured on ALI for at least 30 days (ALI day >30)), we observed acetylated microtubules as a crescent surrounding the apical side of the cell (Fig. 2E, S2D, Movie 7). This microtubule subset, which is not attached to the basal bodies, likely corresponds to the α-tubulin-positive apical microtubule band previously described in mouse trachea epithelial cells (Vladar *et al*., 2012). To study the acetylation pattern of individual microtubules, we measured intensities of acetylated and tyrosinated tubulin along the length of linearized microtubules (Fig. 2F). Apicobasal microtubules showed little acetylation (MT1 in Fig 2E,F), or were acetylated only within the apical part (Fig. 2E,F, MT2). In contrast, crescent microtubules were often fully acetylated (MT3). Quantification of the traced microtubules (the apical 0.34 μm (2%) of the cell were excluded due to very high microtubule density) showed that between ALI day 10 to day >30, acetylated microtubules gradually transitioned from a mostly vertical to a horizontal orientation, in contrast to the majority of microtubules stained for α-tubulin, which had a predominantly vertical orientation (Fig. 2H). In comparison to total (α-tubulin-positive) microtubules, acetylated microtubules or microtubule segments were shorter, especially in mature cultures, and, based on the position of their basal ends, were confined to the apical part of the cell (Fig. 2G,I). To examine microtubule stability, we treated cells with a microtubule-destabilizing drug nocodazole and found that the crescent persisted whereas other parts of the microtubule network depolymerized (Fig. S2E). This is consistent with the notion that the most strongly acetylated microtubule subset, the crescent, is not dynamic.

Since mature MCCs are polarized in the apical plane to coordinate ciliary beating, we hypothesized that the location of the acetylated microtubule crescent would correlate with polarity markers. Proximal planar cell polarity (PCP) protein Vang-like2 (VANGL2) and basal body positioning strongly correlated, consistent polarity corresponding to the oral-lung axis in our model system (Fig. S2F,G). The acetylated microtubule crescent was often located at the cell side opposite to VANGL2 (Fig. S2H,I), suggesting that it correlates with PCP, but no strict correlation between the presence of the apical crescent and basal body alignment was observed (Fig. S2J). To conclude, during MCC maturation, stable microtubules organize into an apical crescent, while the majority of apicobasal microtubules remain dynamic.

### Basal foot nucleates microtubules

As discussed above, apicobasal microtubules are anchored at the basal feet (Fig. 3A,B). To examine whether the basal foot is the site of microtubule nucleation, we imaged microtubule recovery after nocodazole-induced depolymerization (Fig. S3A). Newly polymerized microtubules were visualized by staining for End Binding proteins EB1 and EB3. Immunofluorescence staining showed that after nocodazole washout, basal cells contained one EB1 aster colocalizing with the centrosome (Fig. 3C,D, Fig. S3B, arrow). To examine nucleation events in MCCs, we imaged EB3, as MCCs predominantly express this EB homologue (Fig. S3C)(Schroder et al., 2011). In untreated cells, EB3 localized to basal bodies (Fig. 3C, Fig. S3B-F). In nocodazole-treated cells and directly after nocodazole washout, EB3 strongly colocalized with the tight junction marker zona occludens protein 1 (ZO-1) (Fig. S3F), suggesting that the soluble EB3 pool released by microtubule depolymerization is captured by an apical junction partner. After nocodazole washout, EB3 comets appeared, whereas the junctional EB3 signal faded over time (Fig. S3B,D). In most cells, EB3 comets were initially distributed randomly throughout the cell volume, although they were less abundant in the apical plane; very occasionally, an EB3 aster, originating from a cell-cell junction or in the cytoplasm, was observed (Fig. S3B,D,E). 20 minutes after nocodazole washout, EB3 re-localized to the apical plane in a punctate manner corresponding to the pattern of basal bodies (Fig. S3B,D). Expansion microscopy showed the EB3 puncta colocalize with the basal feet (Fig. 3D). At this resolution, we could observe tyrosinated tubulin, a marker for newly polymerized microtubules, originating from this site. In some instances, we observed a short microtubule with an EB3 comet (Fig. 3D, arrow). These findings indicate that basal foot tips can serve as microtubule nucleation sites, though their activation after nocodazole washout is slow.

**Figure 3:**
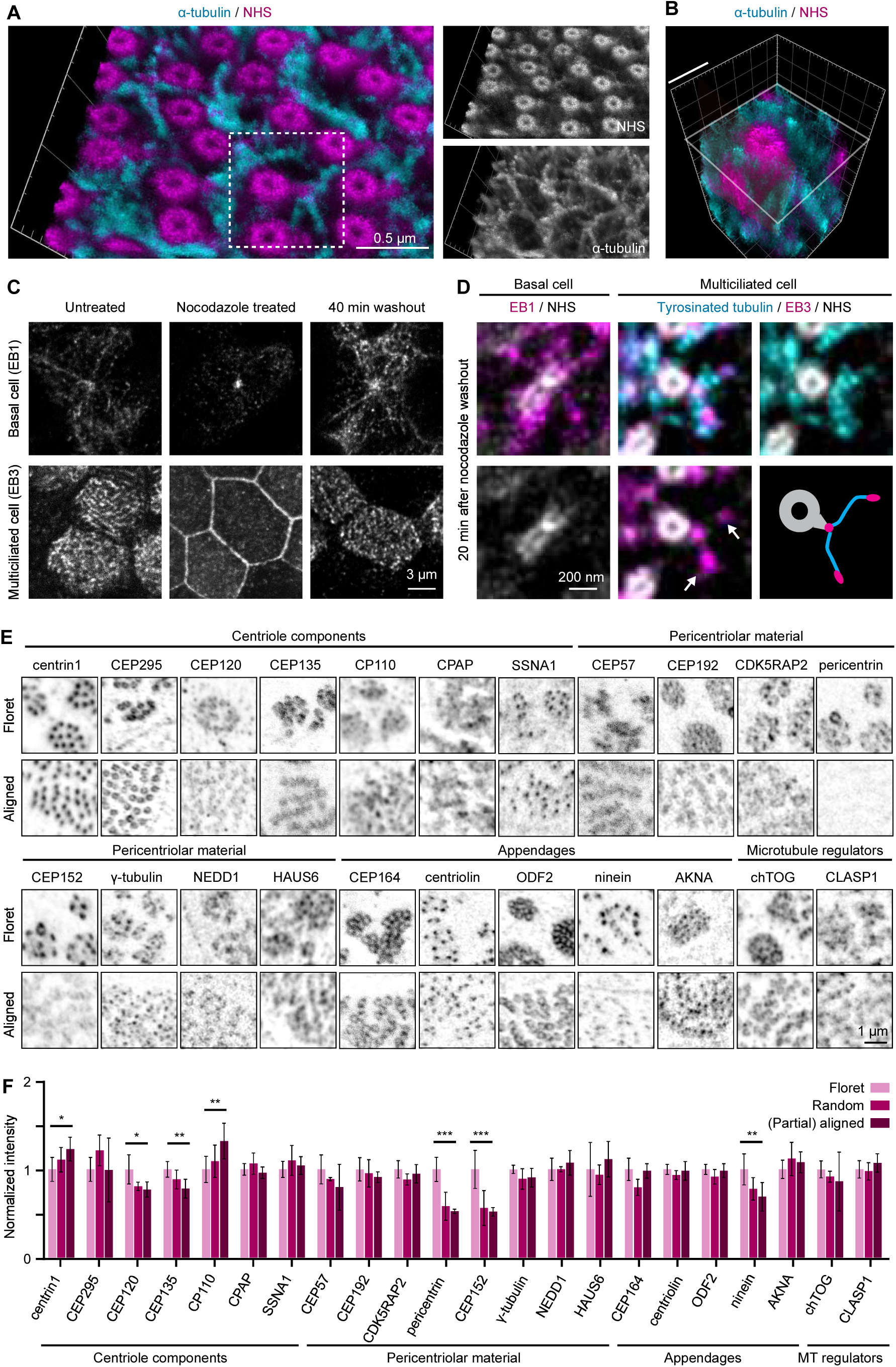
Characterization of the basal body-associated MTOCs in MCCs. (A-B) Volumetric 3D render of a TREx image of an apical region of an MCC stained for α-tubulin (cyan) and NHS-ester (magenta). Box highlights a basal body enlarged in (B). Scale bar in B (corrected for expansion) is ∼200 nm. (C) Microtubule regrowth after nocodazole washout in basal cells and MCCs stained for EB1 and EB3, respectively. (D) Microtubule nucleation after nocodazole washout visualized with TREx in basal cells and MCCs stained for EB1 (magenta) and NHS-ester (white) in basal cells or EB3 (magenta), tyrosinated tubulin (cyan) and NHS-ester (white) in MCCs. White arrows point towards EB3-decorated microtubule ends. (E) Basal bodies in cells with a floret or aligned basal body pattern stained for proteins that localize to the basal body, imaged using STED microscopy. Images represent enlargements of the regions marked by boxes in Fig. S4. (F) Fluorescence intensity (mean±SD; values normalized to floret) of various proteins in cells with a floret, random or (partially) aligned basal body patterns.n, number of analyzed cells: CEP120 and CEP192, n=12; pericentrin and CEP152, n=13; NEDD1, n=15; CEP164, n=16; CEP295, CEP135 and CEP57, n=17; centriolin, n=19; CDK5RAP2, γ-tubulin, ODF2 and ninein, n=20, CP110, AKNA, chTOG and CLASP1, n=21; SSNA1 and centrin1, n=22; CPAP and HAUS6, n=23. Statistical test results are shown only for the comparison between floret and (partially) aligned stage, *P<0.05, **P<0.01, ***P<0.001 is calculated using unpaired non-parametric T test.

### Ninein, CEP120, CEP152 and pericentrin are downregulated during MCC maturation

In order to identify proteins responsible for microtubule nucleation in MCCs, we set out to compare the composition of basal bodies to centrosomes of dividing cells. We categorized proteins as components of the centriole (centrin1, CEP295, CEP120, CEP135, CP110, CPAP and SSNA1), pericentriolar material (PCM) (CEP57, CEP192, CDK5RAP2, pericentrin, CEP152, γ-tubulin, NEDD1 and HAUS6), centriole appendages/basal feet (CEP164, centriolin, ODF2, ninein and AKNA), and microtubule regulators (chTOG, CLASP1) (Fig. 3E and S4)(Camargo Ortega et al., 2019; Conduit et al., 2015; Laporte *et al*., 2024; LeGuennec et al., 2021; Liu et al., 2021; Pfister et al., 2025). Using STED microscopy, we measured protein signal intensity at the basal bodies in MCCs with a floret, random and (partially) aligned basal body patterns at ALI culture day 15 (Fig. 1E,F, 3E,F and S4). The signals of most proteins remained stable during MCC maturation. However, centriole components CEP120 and CEP135 and basal foot protein ninein showed a small but significant reduction during maturation, whereas the signal of centrin1 and CP110 mildly increased. Increased abundance or reactivity of CP110 was surprising, as this protein is the major component of the centiolar cap that is removed during cilia formation (Spektor et al., 2007), and is thought to be downregulated during ciliogenesis (Choksi et al., 2024; Song et al., 2014). Among PCM components, we observed a strong reduction of CEP152, a protein primarily participating in centriole duplication (Cizmecioglu et al., 2010), and pericentrin, a major scaffold for microtubule nucleation in mitosis and interphase (Chen et al., 2022; Doxsey et al., 1994). The presence of PCM proteins at the basal body confirms its role as an MTOC, which, with a few exceptions, shows a relatively constant composition during MCC maturation.

### TREx and computational averaging enable systematic mapping of basal body proteins

To build an accurate 3D sub-micron map localization map of the basal body components, we turned to expansion microscopy. For the ultrastructural cell context, we combined immunostaining of the proteins-of-interest with an NHS ester fluorescent dye labeling (Fig. 2B-C,3A,4C,4G). The NHS signal closely overlapped with that of acetylated tubulin, confirming that it marks the microtubule wall of the basal body (Fig. 4A,B). The well-characterized proteins ODF2 and centriolin localized to the basal body and basal foot, as expected (Fig. 4C). The resolution of TREx was sufficient to resolve the punctate pattern of ODF2 that is characteristic of the nine-fold symmetric structure of the basal body.

**Figure 4:**
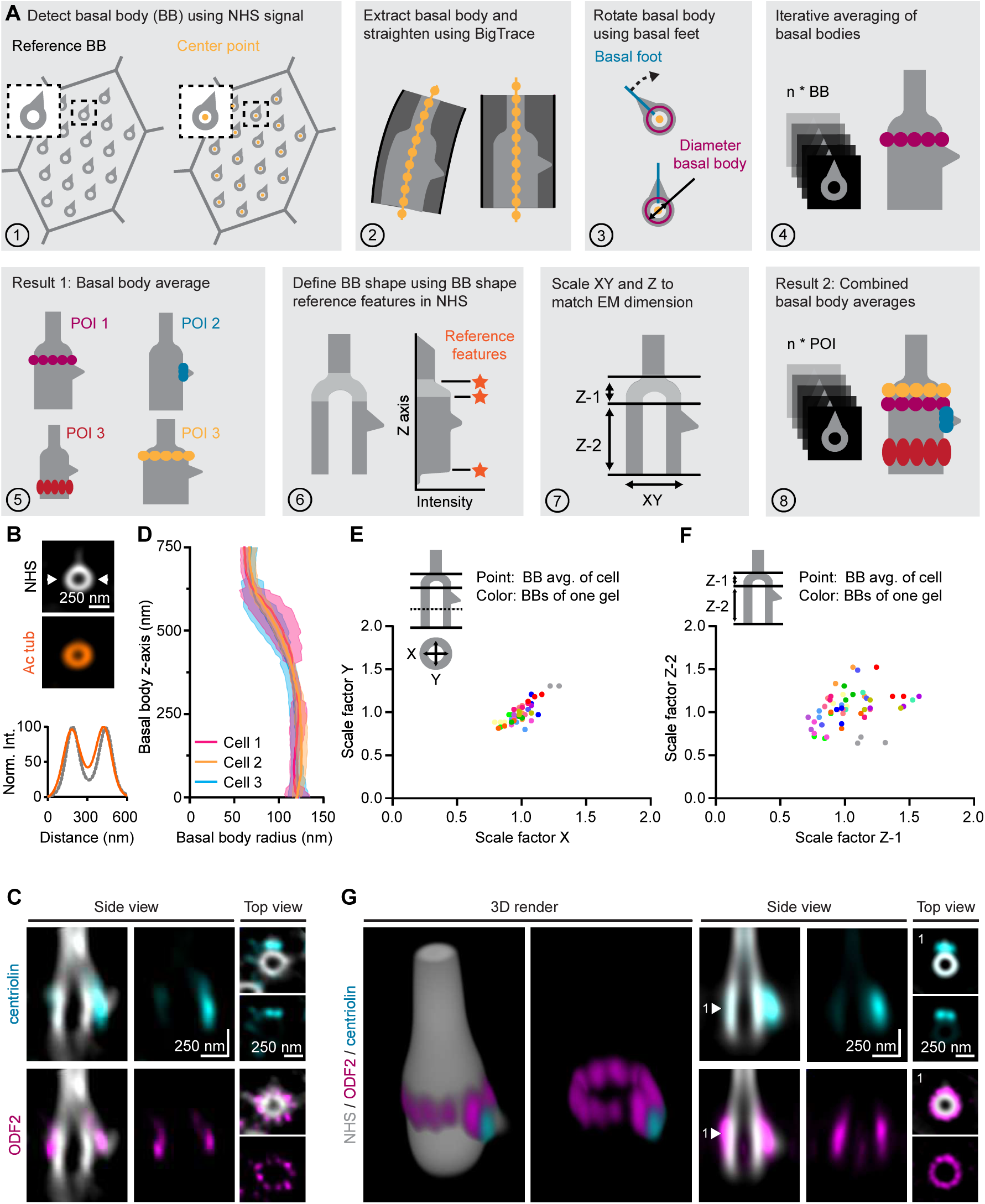
Basal body averaging pipeline and validation. (A) Scheme of the basal body averaging pipeline for cells imaged with TREx, including basal body (BB) detection (1), straightening (2), rotation (3), iterative averaging (4) into basal body averages per protein of interest (POI) (5), detection of basal body shape reference features (asterisks) using NHS-ester intensity profiles (6), scaling in XY and Z dimensions using shape reference features (7) and iterative averaging of basal body averages into the basal body architectural map (8). See Methods for details. ((B) Transverse cross-section of an averaged basal body stained for acetylated tubulin (Ac tub, orange) and NHS-ester (white). Arrows indicate the region of line scan to show the normalized intensity (Norm. Int.) profile of acetylated tubulin (orange) and NHS (gray), shown at the bottom. (C) Longitudinal (left) and transverse cross-section (right) of non-averaged, deconvolved basal body stained for centriolin (cyan), ODF2 (magenta) and NHS-ester (white). (D) Mean±SD of basal body radius along z-axis of three cells acquired from one TREx gel. n, number of analyzed basal bodies: cell 1, n=27; cell 2=45; cell 3=96. (E, F) Scaling factors of basal bodies width in X and Y normalized to the average basal body diameter (E) and basal bodies heights in Z-1 (transition zone) and Z-2 (basal body) segments normalized to the corresponding average lengths (F). Points represent basal body average of one cell. Colors represent basal bodies of cells that were acquired in the same gel. (G) Averaged image of basal body (NHS ester, white) with centriolin (cyan) and ODF2 (magenta) in volumetric 3D render (left) and cross-sections. Numbered arrows in longitudinal cross-section (middle) corresponds to plane of the transverse cross-section (right).

The appearance of the basal bodies often deviated from a round cylinder shape, being squeezed or stretched in arbitrary direction (Fig. 2A, S5A). Compared to relatively immobile centrioles, basal bodies can be deformed by the active motion of cilia, to which they are connected (Junker et al., 2022). This leads to variability of protein positions in individual images. Furthermore, heterogeneity in labeling density, local gel deformations, drift along the z-axis and sample-to-sample variations in the expansion factor can complicate reliable determination of the absolute nanoscale position of individual proteins. An additional complexity arises from the interfering signals of adjacent basal bodies and the basal body tilt with respect to the apical plane, inherent to MCCs. To overcome these challenges, we developed an automated tool to average basal bodies by computationally extracting, aligning and overlaying multiple images (Fig. 4A). In our workflow, the “total protein stain” NHS channel served as a reference 3D shape for the alignment and deformation corrections of protein-of-interest densities. It is an unambiguous 3D template, since its shape is axially asymmetric, and the basal foot determines its unique rotational orientation in the XY plane. This approach avoids an assumption of 9-fold symmetry of centriole-like structures (Chang *et al*., 2023) and allows capture of any asymmetric 3D features (Gaudin et al., 2022).

First, we used the NHS-highlighted shape to automatically detect 70-100 round cross-sections of individual basal bodies from a large field of view covering the apical surface of a single cell. In the next step, for each z-plane, the centerline position of the basal body cross-section was estimated (Fig. 4A-1). Further, the tubular volumes around these centerlines were extracted, straightened (Fig. 4A-2) and rotated in XY plane to align their basal feet (Fig. 4A-3). Next, we performed iterative averaging on 70-100 structures per single cell to obtain the spatial density distribution of proteins-of-interest with respect to the basal body NHS signal (Fig. 4A-4 and Fig. 4A-5). Although the variation in basal body dimensions (both in XY and Z), as measured by NHS signal, was minimal within one cell (Fig. 4D), it increased when basal bodies from cells within one gel or different gels were compared (Fig. 4E, F). This likely reflects cumulative differences in expansion factor, local gel deformations, cell and inter-sample variability. To combine the data from multiple cells and protein stainings together, we scaled each averaged NHS body using their shape and intensity features as reference to a single global average form. The final dimensions of this form were assigned using a reference diameter of basal bodies (250 nm), as measured by EM (Fig. 4A-7)(Li et al., 2012; Vladar et al., 2016). Finally, we registered and combined the homogenized protein averages collected from different experiments (Fig. 4A-8).

**Figure 5:**
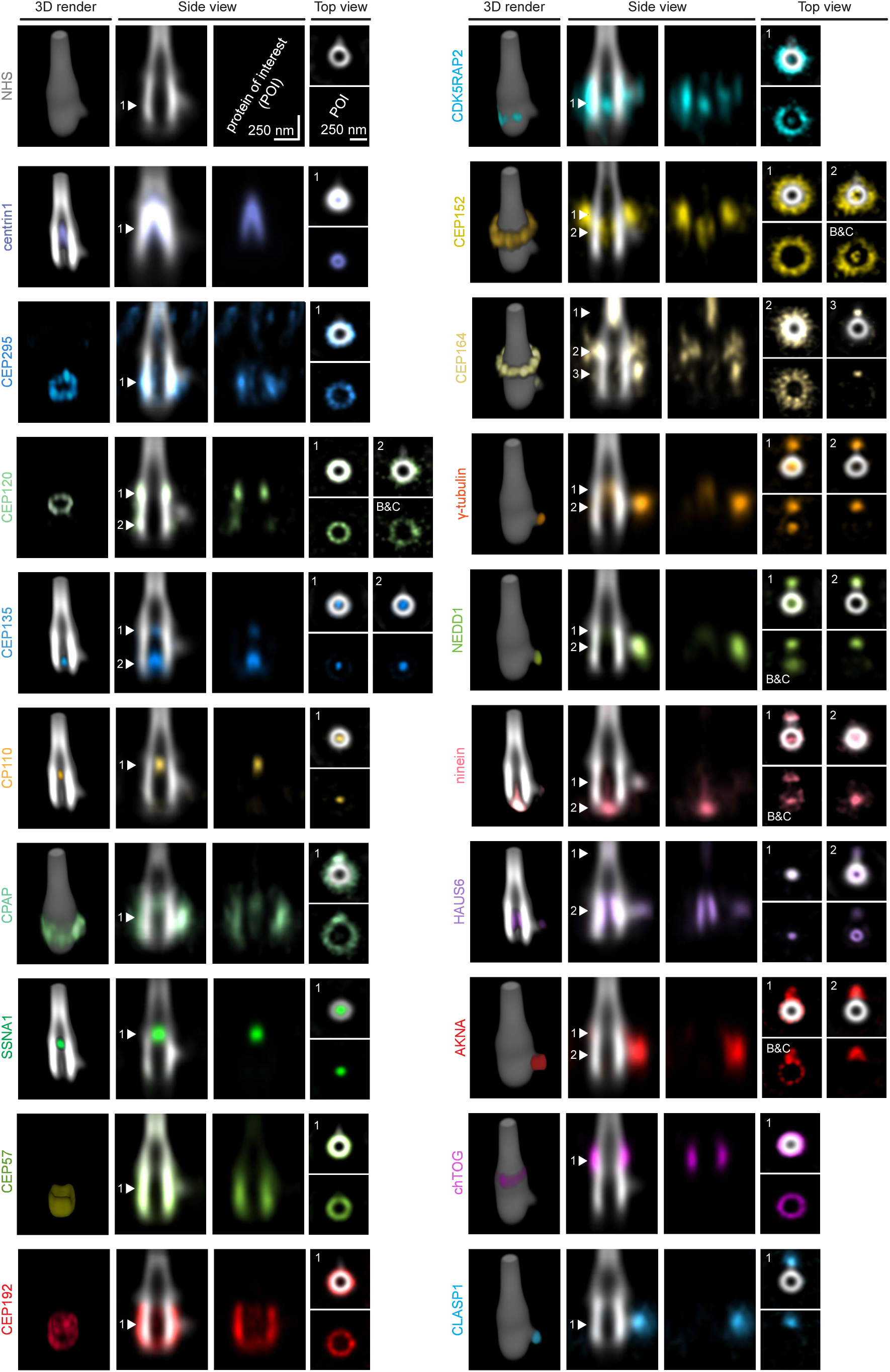
3D architectural protein map of basal bodies in mature MCCs. Gallery of maps for various proteins which localize to the basal body. Colors represent various proteins, with the basal body average (NHS ester) shown in white. For each protein, a volumetric 3D render (left), longitudinal (middle) and transverse cross-section (right) are shown. Numbered arrows in longitudinal cross-section correspond to the plane of the transverse cross-section. For some projection the brightness & contrast (B&C) was changed to show proteins at less prominent positions.

To validate the accuracy of the averaging tool, we focused on centriolin and ODF2 (Fig. 4G). Centriolin localizes to the basal foot as two distinct puncta adjacent to the base of the basal foot, resembling that of basal foot protein CEP112 (Nguyen *et al*., 2020). ODF2 forms a ring around the basal body, and staining densities of individual basal bodies showed clear nine-fold symmetry patterns with the expected 40° angle between puncta, in line with published data (Fig. S5A,B)(Chang *et al*., 2023; Chong *et al*., 2020; Ishikawa *et al*., 2005). Apart from the ring pattern, the average 3D map revealed an additional local accumulation of ODF2 in the vicinity of the basal foot. It is possible that ODF2 binds at two different sites (Fig. 4G), in line with recent observations showing that ODF2 is present both at distal and subdistal centriole appendages (Chang *et al*., 2023).

Based on the average protein distributions, we built a joint architectural 3D map of the basal body (Fig. 5, Movie 8). From the projection of the spatial protein densities on the center-to-periphery axis, protein patterns were categorized into three distinct groups: luminal, ring-like and basal foot localization (Fig. 5 and 6A). The corresponding projection on the longitudinal axis showed a stacked organization of the ring-like and luminal proteins distributed from the proximal to the distal end (Fig. 6B). As discussed below, the patterns of some proteins were similar to those in centrioles, whereas others were basal body-specific (Fig. 6C).

**Figure 6:**
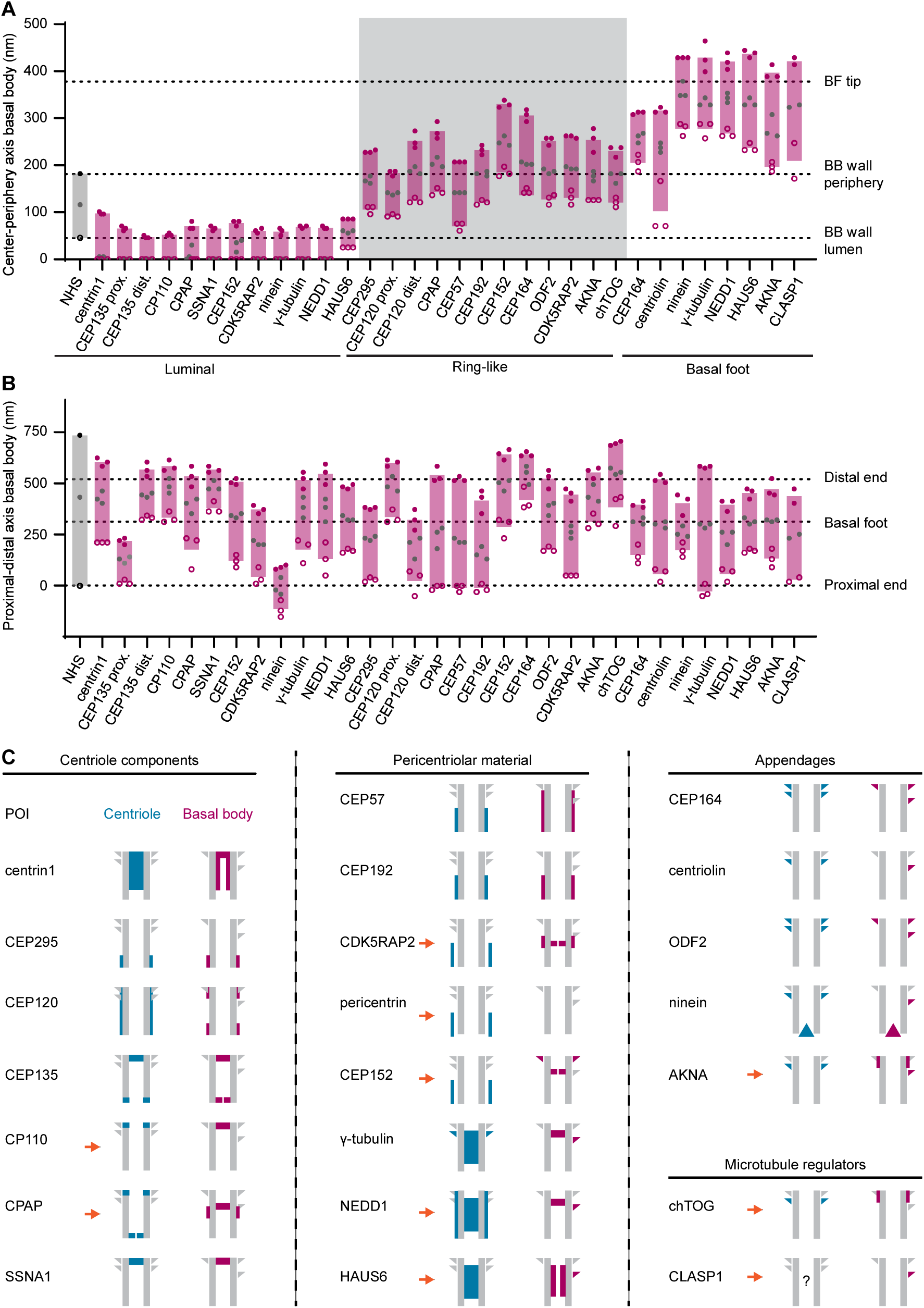
Quantification of protein localization in basal bodies and their comparison with centrioles. (A) Position of various proteins along the radius relative to the averaged basal body visualized by NHS ester (gray), grouped by localization: luminal, ring-like and basal foot. Solid gray circles represent maximum fluorescence intensity of protein. Solid and hollow magenta circles (black for NHS) represent the position of half maximum fluorescence intensity of protein. Bars represent coverage of protein based on full width at half maximum intensity. Dashed lines show the positions of the basal foot (BF) tip and the luminal or peripheral border of the basal body (BB) wall. Abbreviations: proximal protein position (prox.) and distal protein position (dist.). n, number of analyzed basal body averages: NHS, n=1 (combined basal body average); proteins, n=3 (from three different cells). (B) Longitudinal position of various proteins relative to the averaged basal body visualized by NHS-ester (gray) grouped and plotted in the same way as in panel A. (C) Schematic comparison of protein positions between centrioles (blue, published work) and basal bodies (magenta, this study). Orange arrows point towards proteins with different localization in the two structures. Position in centrioles is based on: centrin1 (Laporte *et al*., 2024; Schweizer *et al*., 2021); CEP295 (Laporte *et al*., 2024; Tsuchiya *et al*., 2016); CEP120 (Mahjoub *et al*., 2010; Tsai *et al*., 2019); CEP135 (Laporte *et al*., 2024); CP110 (Laporte *et al*., 2024; Spektor *et al*., 2007); CPAP (Iyer *et al*., 2025; Laporte *et al*., 2024); SSNA1 (Huang *et al*., 2025); CEP57 (Watanabe *et al*., 2019); CEP192, CDK5RAP2, pericentrin, NEDD1 and HAUS6 (Schweizer *et al*., 2021); CEP152 (Laporte *et al*., 2024; Sullenberger *et al*., 2023); γ-tubulin (Chang *et al*., 2023; Laporte *et al*., 2024; Schweizer *et al*., 2021); CEP164 (Laporte *et al*., 2024; Yang *et al*., 2018); ODF2 (Chang *et al*., 2023; Chong *et al*., 2020); ninein (Chang *et al*., 2023); AKNA (Camargo Ortega *et al*., 2019); chTOG (Ali et al., 2023).

**Figure 7:**
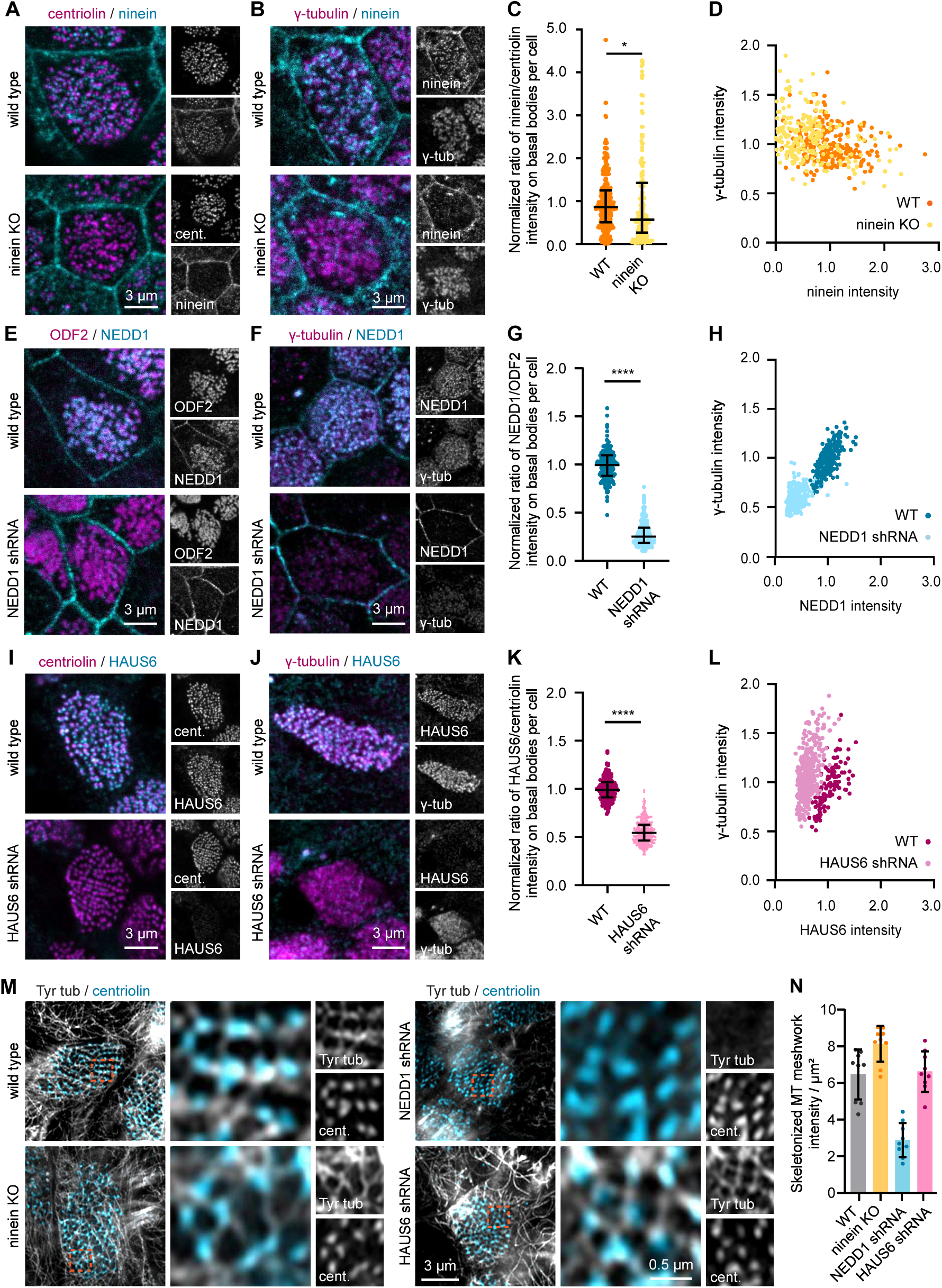
Functional analysis of basal foot components in MCCs. (A, B) Basal bodies of wild type and ninein knockout (KO) cells stained for ninein (cyan) and centriolin (A, magenta) or γ-tubulin (B, magenta). (C) Normalized ratio of fluorescence intensity (median±IQR) of ninein to centriolin staining at basal bodies of the same wild type and ninein KO MCCs. Each point represents one MCC; number of analyzed cells, from 3 independent experiments: wild type, n=201; ninein KO, n=147. (D) Fluorescence intensity of ninein versus γ-tubulin intensity at basal bodies in wild type and ninein KO cells. All datapoints are normalized to the normalized to 1, wild type average. Each point represents one MCC; n, number of analyzed cells from 3 independent experiments: wild type, n=230; ninein KO, n=206. (E, F) Basal bodies of wild type and NEDD1 depleted cells stained for NEDD1 (cyan) and ODF2 (E, magenta) or γ-tubulin (F, magenta). (G) Normalized ratio of fluorescence intensity (median±IQR) of NEDD1 to ODF2 staining at basal bodies in wild type and NEDD1 depleted MCCs, plotted as in panel C. n, number of analyzed cells, from 3 independent experiments: wild type, n=173; NEDD1 shRNA, n=261. (H) Fluorescence intensity of NEDD1 versus γ-tubulin intensity at basal bodies in wild type and NEDD1 depleted MCCs, plotted as in panel D. n, number of analyzed cells, from 3 independent experiments: wild type, n=272; NEDD1 shRNA, n=267. (I, J) Basal bodies of wild type and HAUS6 depleted cells stained for HAUS6 (cyan) and centriolin (I, magenta) or γ-tubulin (J, magenta). (K) Normalized ratio of fluorescence intensity (median±IQR) of HAUS6 to centriolin staining at basal bodies in wild type and HAUS6 depleted MCCs, plotted as in panel C. n, number of analyzed cells, from 3 independent experiments: wild type, n=117; HAUS6 shRNA, n=289. (L) Fluorescence intensity of HAUS6 versus γ-tubulin intensity at basal bodies of wild type and HAUS6 depleted cells, plotted as in panel D. n, number of analyzed cells, from 3 independent experiments: wild type, n=116; HAUS6 shRNA, n=378. (M) Apical microtubule meshwork of MCCs stained for tubulin (white) and NHS-ester (cyan) and imaged using Airyscan2. Orange box marks region enlarged in zoom. Comparison between wild type, ninein KO, NEDD1 shRNA and HAUS6 shRNA cells. (N) Quantification of microtubule apical meshwork using tubeness filter and subsequent thresholding and skeletonization. Comparison between wild type, ninein KO, NEDD1 shRNA and HAUS6 shRNA cells. n, number of analyzed cells: wild type, n=9; ninein KO, n=9; NEDD1 shRNA, n=9; HAUS6 shRNA, n=9. For all bar graphs in this image, ****P<0.001 is calculated using unpaired two-tailed Mann-Whitney U test.

### Comparison of the localization of centriole and basal body components

In line with previous work on the centriole (Laporte *et al*., 2024), centrin1 was found in the basal body lumen, where it formed a cone-shaped accumulation, starting as a ring at the proximal end and tapering more distally (Fig. 5, 6). CEP295 formed a ring around with the proximal part of the basal body wall, in line with its positioning in centrioles (Izquierdo et al., 2014; Laporte *et al*., 2024). CEP120, which binds along the centriole wall (Tsai et al., 2019) and is known to be upregulated during MCC development (Mahjoub et al., 2010), localized as two distinct rings at the proximal and distal end of the basal body with a diameter close to that of the microtubule wall. We detected a clear nine-fold symmetry for the distal CEP120 ring, which appeared brighter than the proximal ring (Fig. S5A,B).

Another conserved microtubule-binding centriole assembly factor, CEP135, which localizes to the centriole wall at the proximal end and to the centriole lumen at the distal end (Laporte *et al*., 2024), was positioned in the lumen at both ends of the basal body (Fig. 5, 6). In centrioles, the proximal pool of CEP135 corresponds to the site where cartwheel proteins like SAS-6 are found (Laporte *et al*., 2024). In cells with motile cilia, the homologue of CEP135, Bld10, participates in the formation of the central pair of microtubules (Carvalho-Santos et al., 2012) and helps to maintain the structural integrity of the basal body during ciliary beating (Bayless et al., 2012). Localization of CEP135 is consistent with these functions.

One major difference between centrioles and basal bodies is the presence of the distal centriolar cap containing CP110 and CPAP (Fig. 5,6, S5 and S6A), which control microtubule plus end elongation (Iyer et al., 2025; Spektor *et al*., 2007). In centrioles, CPAP localizes to the proximal end, the distal cap and the PCM (Iyer *et al*., 2025; Laporte *et al*., 2024; Vasquez-Limeta et al., 2022), whereas in basal bodies, CPAP formed a diffuse ring around the basal body with a significant enrichment at the basal foot. CP110 blocks centriolar microtubule plus ends and must be removed to allow formation of ciliary doublets (Choksi *et al*., 2024; Song *et al*., 2014; Spektor *et al*., 2007). Surprisingly, CP110 did not disappear from basal bodies, but localized as a small sphere in the distal basal body lumen. In close proximity to luminal CP110 was SSNA1, a protein recently shown to organize the distal luminal centriolar network (Huang et al., 2025; Pfister *et al*., 2025). Together with the observation that CP110 intensity increased when MCCs transitioned from a floret to an aligned basal body pattern (Fig. 3E,F), this suggests that CP110 is not removed but rather displaced from the microtubule wall to the lumen.

### Several major PCM proteins do not accumulate at the basal foot

PCM organization at the basal body also showed similarities and differences from that of centrosomes in dividing cells (Fig. 5,6, S5 and S6A). CEP57, a PCM organizing protein (Watanabe et al., 2019), and CEP192 encircled the basal body, with enrichment towards its proximal end, at the same region as CEP295, to which CEP192 is known to bind (Tsuchiya et al., 2016). CDK5RAP2, which in centrosomes colocalizes with CEP57 and CEP192 at the proximal centriole end (Schweizer et al., 2021), was present along the longitudinal axis of the basal body in a non-symmetrical manner. Two other major PCM components, CEP152 and pericentrin, were downregulated during MCC maturation. The intensity of pericentrin, which participates in organizing the PCM layers around centrioles (Mennella et al., 2012), decreased to very low levels in mature MCCs (Fig. 3E,F). CEP152 formed a ring with a nine-fold symmetry in the distal appendage region of the basal body in contrast to its proximal localization on the centriole, where CEP152 is part of the platform for procentriole formation (Fig. S6A)(Laporte *et al*., 2024; Sullenberger et al., 2023). The loss of pericentrin in mature MCCs and the ring-like localization of other PCM components discussed above suggest that they are not responsible for concentrating MTOC activity at the tip of the basal foot (Fig. S6A).

### Localization of appendage proteins provides candidates for MTOC activity

Positioning of centrioles and basal bodies critically depends on their appendages. Distal appendages are required for docking of cilia and vesicles to the plasma membrane (Tanos et al., 2013). Their component CEP164 (Schmidt et al., 2012) formed a large distally located ring with a clear nine-fold symmetry (Fig. 5,6, S5 an S6A), very similar to its localization at the centriole (Laporte *et al*., 2024). Furthermore, CEP164-positive signal was also observed inside the cilium and on the basal foot.

Basal feet, the subdistal appendages of basal bodies, consist of several conserved components (Nguyen *et al*., 2020), such as ODF2 and centriolin, discussed above (Fig. S6A). The tip of the basal foot accumulates γ-TuRC and its binding partner NEDD1, in agreement with the fact that it serves as a MTOC (Clare *et al*., 2014; Nguyen *et al*., 2020). We confirmed this observation and showed that, similar to centrioles (Schweizer *et al*., 2021), a pool of γ-TuRC and NEDD1 is also present in the basal body lumen (Fig. 5 and 6). In centrioles, luminal γ-TuRC was proposed to enhance centriole stability or to be sequestered for release during mitotic onset to participate in spindle formation (Gao et al., 2025; Schweizer *et al*., 2021). The former function might also apply to the luminal γ-TuRC in the basal body. CDK5RAP2, an activator of γ-TuRC (Choi et al., 2010), was also present in basal body lumen (Fig. 5), but was not enriched at the basal foot.

Another potential candidate for promoting microtubule nucleation and anchoring is ninein, an abundant basal foot component (Delgehyr et al., 2005; Nguyen *et al*., 2020). We confirmed ninein localization to the peripheral part of the basal foot (Fig. 5). The precise localization of ninein varied between basal bodies (Fig. S6B), suggesting that its position is not controlled tightly. A pool of ninein was also present at the most proximal end of the basal body (Goldspink *et al*., 2017; Mogensen et al., 2000), where it located ∼120 nm outside of the basal body structure in a plug-like manner. In centrioles, the proximal ninein pool participates in the formation of the centrosome linker (Dang and Schiebel, 2022), but its function in the basal body is unclear.

The presence of NEDD1 together with γ-TuRC prompted us to investigate the localization of their binding partner Augmin/HAUS, a protein complex responsible for branching microtubule nucleation (Petry et al., 2013; Zhang et al., 2022). The HAUS6 subunit of the complex was localized to the basal body lumen, similar to centriolar lumen (Gao *et al*., 2025; Schweizer *et al*., 2021). The luminal pool of HAUS6 was structured as a hollow cylinder, while γ-TuRC and NEDD1 were visible as a blob, in line with the idea that HAUS tethers γ-TuRC to the inner centriole wall (Gao *et al*., 2025; Schweizer *et al*., 2021). Surprisingly, a substantial HAUS6 signal was present at the basal foot (Fig. 5), suggesting that it might participate in the MTOC function.

We also examined the localization of AKNA, a recently identified component of subdistal appendages, required for γ-TuRC-dependent microtubule nucleation in neural stem cells (Camargo Ortega *et al*., 2019). AKNA strongly localized to the peripheral end of the basal foot and formed a weak ring close to that of ODF2 (Fig. 5). AKNA was shown to be able to recruit microtubule polymerase chTOG, which promotes microtubule nucleation (Camargo Ortega *et al*., 2019; Thawani et al., 2018). However, chTOG did not accumulate at the basal foot and instead surrounded the distal basal body, at a position similar to the AKNA ring, above the basal foot (Fig. 5). The puncta of both chTOG and AKNA located around the distal end of the basal body showed nine-fold symmetry (Fig. S5A,B). Finally, we examined the position of another factor promoting γ-TuRC-dependent microtubule nucleation, CLASP (Rai et al., 2024), and found that CLASP1 was strongly enriched at the basal foot tip (Fig. 5). Overall, the basal body map identified a distinct set of proteins to be tested as functional components of the basal body MTOC.

### γ-TuRC recruitment to the basal foot depends on NEDD1, but not ninein or Augmin

To identify proteins required for γ-TuRC recruitment to the basal foot, we systematically depleted γ-TuRC-targeting factors present at this site. To be able to perform multiple rounds of selection, validation and differentiation of genome-edited cell lines, we made use of immortalized human airway cells that we have recently generated (van Grinsven et al, in preparation).

Our best candidate was ninein, as it has been strongly implicated in microtubule nucleation and anchoring at centriolar subdistal appendages as well as other sites (Delgehyr *et al*., 2005; Goldspink *et al*., 2017; van Grinsven and Akhmanova, 2025). We depleted ninein by CRISPR/Cas9 mediated knockout using three gRNAs targeting exon 5 (Fig. S7A). Validation in basal cells using Western blot showed two ninein isoforms in airway cells which were decreased in the edited cells (Fig. S7B). The heterogeneity in the ninein knockout cell population was confirmed by immunofluorescence staining and genomic sequencing (Fig. S7C,D,F). Next, we studied the effects of ninein loss in the MCCs of the airway on a single cell level (Fig. S7E). To take into account the variation in basal body number per cell, we normalized the intensity of ninein to centriolin, a basal body marker (Fig. 7A,C). Although ninein intensity at the basal body decreased, this had no impact on γ-tubulin intensity (Fig. 7B,D). Furthermore, inspection of the apical microtubule network by Airyscan super-resolution microscopy showed that microtubules were still anchored to basal feet in ninein knockout cells, as shown by microtubule overlap with centriolin and the presence of apical microtubule meshwork (Fig. 7M,N). Moreover, microtubule network reestablished normally after nocodazole washout (Fig. S7I). Thus, ninein does not appear essential for γ-TuRC-dependent microtubule organization at the basal foot.

Since γ-TuRC recruitment to the centrosome in cycling cells is mediated by adaptor protein NEDD1 (Haren et al., 2006; Luders et al., 2006), we next studied MCCs depleted of NEDD1. NEDD1 is required for cell division (Haren *et al*., 2006; Luders *et al*., 2006), and therefore could not be stably knocked out; instead, we designed doxycycline inducible shRNA constructs against NEDD1 (Fig. S7G). We induced the expression of NEDD1 shRNA at the onset of MCC differentiation for a period of 30 days and successfully depleted NEDD1 from the basal bodies, as determined by normalizing NEDD1 signal to that of ODF2 (Fig. 7E,G). In line with its role as a γ-TuRC adaptor protein, NEDD1 depletion significantly reduced γ-tubulin intensity at the basal bodies (Fig. 7F,H). The apical microtubule network typically associated with the basal bodies was partially absent, and microtubules that remained associated with the basal feet or cell-cell junctions (Fig. 7N,M, S7J). Moreover, NEDD1-depleted cells showed reduced EB3 accumulation around basal bodies after nocodazole washout (Fig. S7I), suggesting a nucleation defect, but their basal bodies were still aligned (Fig. S7K).

We also generated a doxycycline-inducible shRNA construct against HAUS6 (Fig. S7H). Long-term depletion in MCCs significantly reduced the HAUS6 intensity at the basal bodies, but this did not lead to a reduction in γ-tubulin levels (Fig. 7I-L). Therefore, HAUS complex is not essential for γ-TuRC targeting to the basal foot. In line with these data, the apical microtubule network and microtubule nucleation after nocodazole washout were unchanged in HAUS6-depleted cells compared to the untreated cells (Fig. 7M,N, S7I). To conclude, among the tested adaptor proteins, only NEDD1 turned out to be essential for γ-TuRC-dependent microtubule organization at the basal foot.

## Discussion

In this study, we visualized microtubule organization and basal body architecture in airway epithelial cells using expansion microscopy combined with computational analysis. We confirmed that epithelial differentiation is associated with a shift to non-centrosomal microtubule networks. In secretory cells, microtubule organization appears random, while in MCCs, basal foot tips nucleate and anchor most apical and apicobasal microtubules. In mature MCCs, these microtubules are mostly short-lived, as they are sensitive to nocodazole and are not particularly enriched in acetylated or detyrosinated tubulin. The dynamic nature of these microtubules correlates with the fact that they are dependent on the NEDD1-γ-TuRC complex, reminiscent of spindle microtubules that are also known to be dynamic (Haren et al., 2006; Luders et al., 2006).

In addition to dynamic microtubules, mature MCCs contain a distinct population of stable, acetylated microtubules: a horizontally positioned crescent with the minus-ends detached from the basal feet. How this microtubule structure is formed is unclear, but this could involve specific microtubule-stabilizing proteins such as CLAMP/Spef1 (Werner et al., 2014) or CAMSAP3 (Kimura et al., 2021; Robinson et al., 2020; Saito et al., 2021; Usami *et al*., 2021). The few vertically positioned acetylated microtubules present in mature MCCs initiate at the apical side and often have a sharply delineated non-acetylated distal segment, which is likely formed when a long-lived, acetylated microtubule starts to depolymerize, is rescued and regrows. The presence of a distinct subset of acetylated microtubules suggests that the recently proposed model that damage caused by stepping kinesin-1 induces an exponential gradient of microtubule acetylation (Andreu-Carbo et al., 2024) may not apply to MCCs.

To get functional insight into the MTOCs in MCCs, we mapped the components of basal bodies by computational averaging. As expected, basal bodies display similarities with centrioles of cycling cells: some centriole wall components such as CEP120 and CEP295, luminal proteins including centrin1 and SSNA1, and the luminal populations of HAUS, NEDD1 and γ-TuRC show comparable localization within the two structures. Other structural components localize differently: for example, on basal bodies the proteins capping the distal centriole end, CP110 and CPAP, enrich in the lumen and on the outer surface, respectively. Another differentially localized protein is CEP152, which forms a platform for centriole duplication at the proximal centriole end but is localized distally and in the lumen of basal bodies. Whether these localization patterns are associated with specific functions of CP110, CPAP and CEP152 in ciliated cells, or merely reflect their affinity for other, functionally important basal body components in conditions when their centriolar function is not needed, is unclear.

The components of distal and subdistal appendages, CEP164, ODF2, centriolin, ninein and AKNA, localize similarly in centrioles and basal bodies, with the obvious difference that in centrioles both types of appendages display a nine-fold symmetry, whereas in basal bodies, there is only one subdistal appendage, the basal foot. The latter structure was a major focus of our study, because in MCCs it represents the main site of γ-TuRC-dependent microtubule organization. This feature is unique for MCCs, because in centrosomes, microtubules emerge not only from the subdistal appendages but also from the γ-TuRC-containing PCM associated with the centriole wall (Chong *et al*., 2020; Mogensen *et al*., 2000). In spite of this architectural difference, some PCM components, such as CEP57 and CEP192, retain their localization around the basal body cylinder, while others, such as CPAP, enrich at the basal foot. Since γ-TuRC and microtubules are not attached to the proximal end of the basal body, CEP57 and CEP192 do not seem to participate in microtubule organization. Pericentrin, another major PCM component, disappears from basal feet of mature MCCs altogether.

While we demonstrate a role for NEDD1 in microtubule nucleation on the basal foot, the mechanism of NEDD1-γ-TuRC tethering and γ-TuRC activation at this site remains unclear. The HAUS complex, with its unexpected accumulation at the basal foot, appeared to be an excellent candidate, but the results of depletion of HAUS6 argue against its essential role within the basal foot MTOC. Furthermore, we could find no support for an essential function of another microtubule-anchoring centrosome and basal foot component, ninein (Nguyen *et al*., 2020), though it is possible that ninein and HAUS operate in a redundant manner. The role of yet another potential player, AKNA, known from a study on subdistal appendages in neural stem cells (Camargo Ortega *et al*., 2019), still needs to be investigated. Among the known activators of γ-TuRC (Choi *et al*., 2010; Rai *et al*., 2024; Thawani *et al*., 2018), only CLASP1 was specifically enriched at the basal foot, and its function, potentially redundant with CLASP2, would need to be analyzed. Two other major activators of γ-TuRC,

CDK5RAP2 and chTOG, are distributed in a ring-like pattern that shows only limited overlap with the basal foot but could still participate in microtubule nucleation.

Taken together, we showed that microtubule organization dramatically changes during differentiation of airway epithelium. Fully differentiated MCCs organize their microtubules around the basal feet. Whereas in cells with a radial microtubule system, one or two centrosomes must be able to populate the whole cell with microtubules, a ciliated cell has numerous (∼70-100) basal bodies. This explains why, compared to centrosomes, the activity of each basal foot associated MTOC is quite low, with only a few microtubules attached to each basal body. This function is supported by the unique composition of basal body MTOC, which represents a combination of building blocks present in other mammalian MTOCs of dividing and differentiated mammalian cells.

## Author contributions

E.J.G. conceptualized, wrote, visualized, performed and analyzed all the experiments in the paper with help of K.B.A, who performed and analyzed parts of Fig. 3E, 3F and Fig. S3B. E.A.K. conceptualized and developed software for microtubule tracing and basal body averaging and wrote the manuscript. F.C. generated the RPE-1 ninein knockout cell line. J.M.B. and L.C.K. conceptualized, reviewed and edited the manuscript, and acquired funding. A.A. supervised the study, conceptualized and wrote the manuscript, and acquired funding.

## Acknowledgement

This work was supported by the Netherlands Organization for Scientific Research (NWO) Gravitation programme IMAGINE! (project number 24.005.009). F.C. was supported by China Scholarship Council. We thank Jens Lüders (Institute for Research in Biomedicine, Spain), Jen-Hsuan Wei (Institute of Molecular Biology Academia Sinica, Taiwan), Magdalena Götz (Institute of Stem Cell Research, Germany) and Lynne Cassimeris (Lehigh University, USA) for the gift of materials, Nefeli Itakisiou for help with cell culture and Jip Manshande for help with optimization of experiments.

## Legends to Supplemental Figures

**Figure S1, related to Figure S1.**
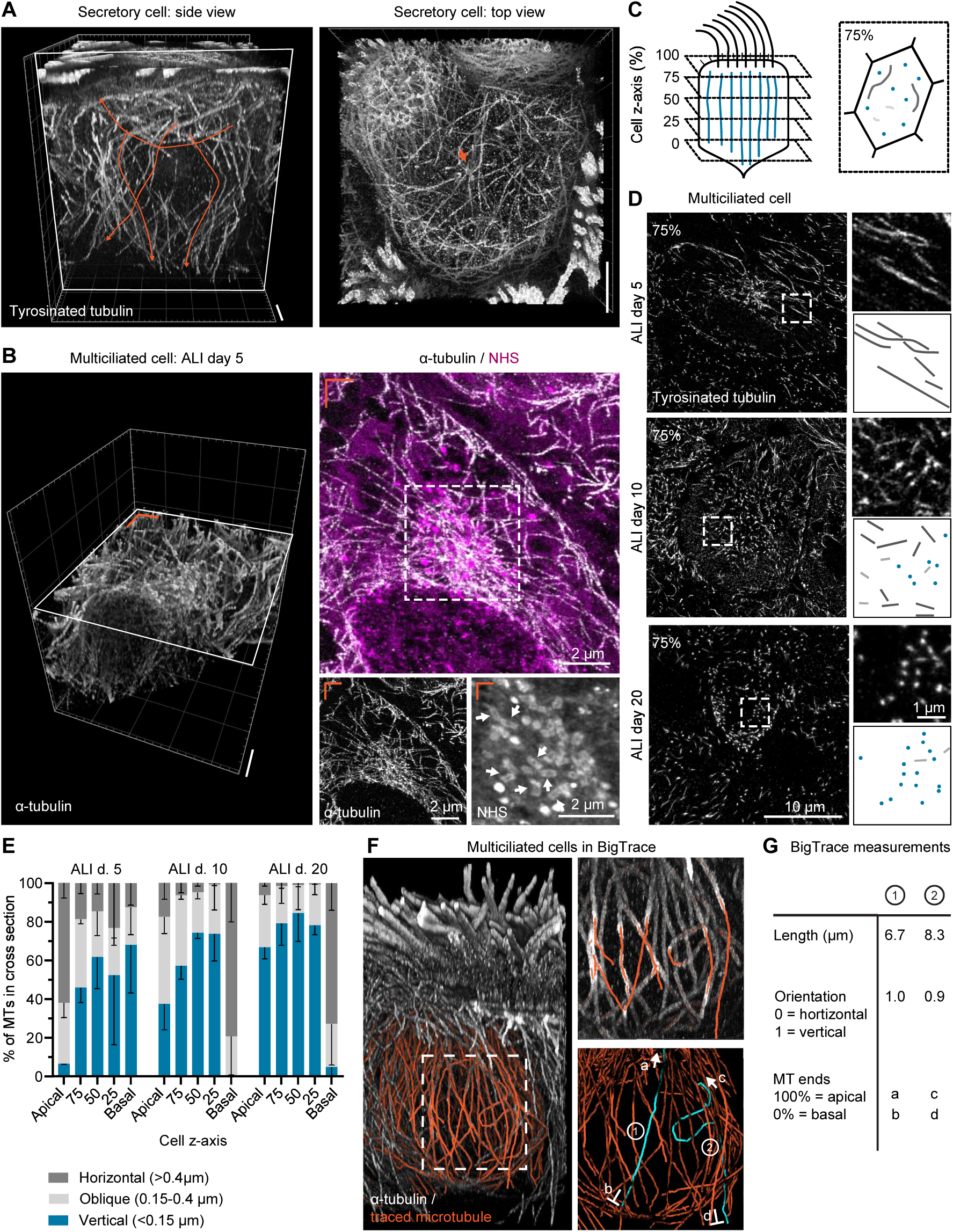
Microtubule network of MCCs and its quantification. (A) Volumetric 3D render of a TREx image of microtubule network in secretory cell stained for tyrosinated tubulin (white); side view and top view (corresponds to Fig. 1H). Orange microtubule traces show variation in microtubule orientation (left) and an orange arrowhead highlights clustered microtubule ends (right). Scale bar (corrected for expansion) is ∼2μm. (B) Volumetric 3D render of a TREx image of microtubule network in MCC at ALI day 5 stained for α-tubulin (white), and a maximum intensity projection of the cross-section of an overlay with NHS ester staining (magenta) (right) (corresponds to Fig. 1I). White box marks the region enlarged in zoom. White arrows in the NHS channel indicate basal bodies. Orange lines in left corner indicate the orientation of maximum intensity projection in relation to 3D render. Scale bar (corrected for expansion) is ∼2μm. (C) Scheme of microtubule density map. The cell axis was divided in 5 cross-sections, where 0% is basal side and 100% is apical side. In each cross-section, the number of microtubule (corrected for cell area) and angle (horizontal = dark gray, oblique = light gray, vertical = blue) is counted. (D) Examples of cell cross-sections at 75% of MCCs at ALI day 5, 10 and 20. White box marks region enlarged in zoom. Scheme indicates how microtubules were scored (see panel C). (E) Percentage of microtubules with different orientation: horizontal (>0.4 μm), oblique (0.15-0.4 μm) or vertical (<0.15 μm) in microtubule density map as shown in Fig. S1B; comparison between MCCs at ALI day 5, 10, and 20. n, number of analyzed cells: ALI day 5, 10 and 20, n=3. (F) 3D tracing of microtubules using FIJI BigTrace plugin. 3D render shows volumetric data (gray), traced microtubules (orange) and selected traced microtubules 1 and 2 (cyan). (G) Example of quantitative measurements of highlighted microtubules 1 and 2 in (F) using BigTrace plugin.

**Supplemental Figure S2, related to Figure 2.**
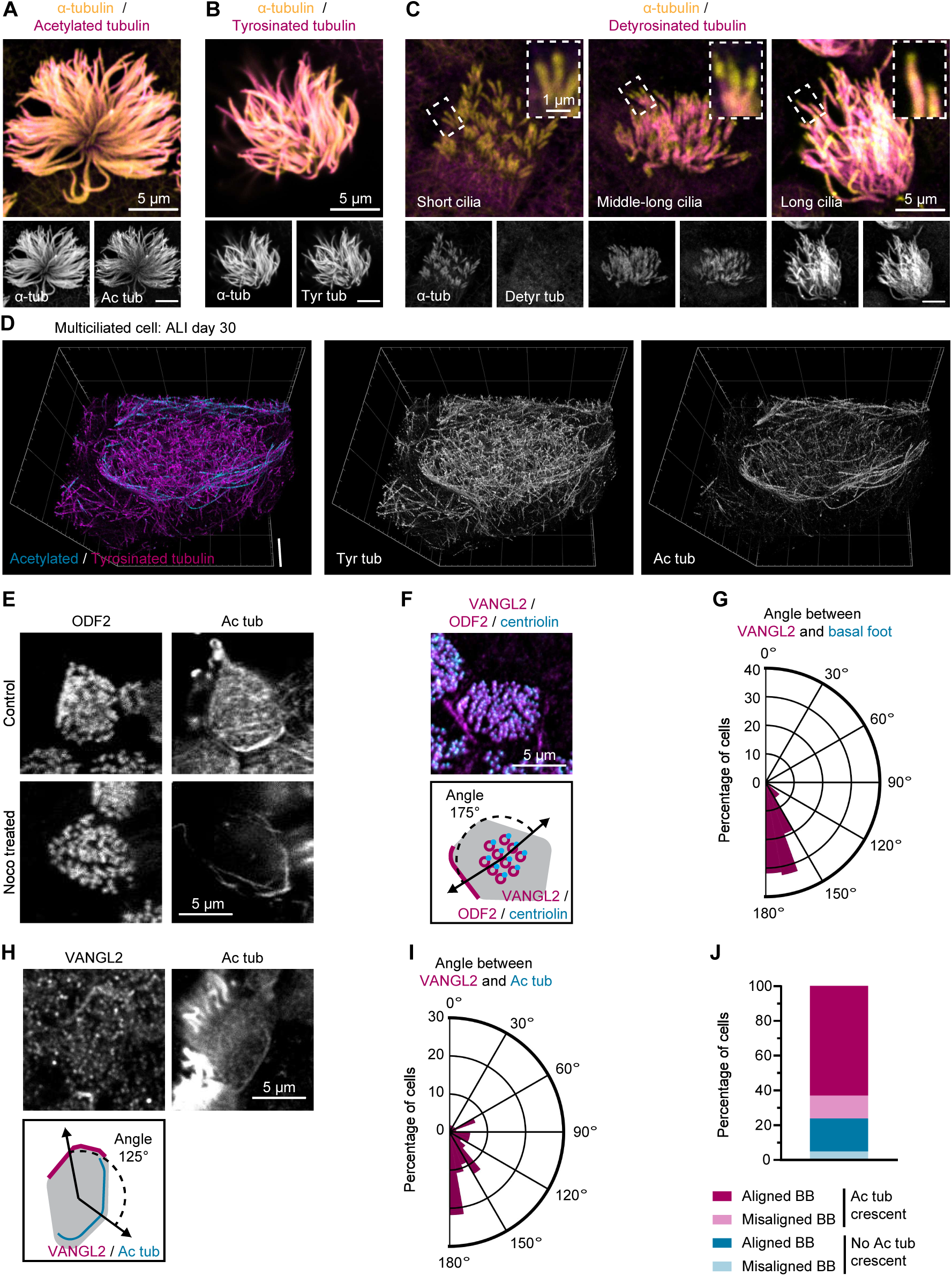
Visualization of post-translationally modified microtubule subsets in MCCs. (A-B) Cilia from MCCs at ALI day 20 stained for α-tubulin (yellow) and acetylated tubulin (A, magenta) or tyrosinated tubulin (B, magenta). (C) Cilia during elongation (short cilia, middle-long cilia and long cilia represents left-to-right) stained for α-tubulin (yellow) and detyrosinated tubulin (magenta). Box highlights ciliary tips enlarged in the inset. (D) Top view volumetric 3D render of TREx image of acetylated microtubule crescent of MCCs at ALI day 30 (corresponding to 3D render in Fig, 2E) stained for acetylated (cyan) and tyrosinated (magenta) tubulin. Scale bar (corrected for expansion) is ∼2μm. (E) Apical acetylated microtubule crescent in control or nocodazole treated MCCs stained for ODF2 (gray) and acetylated tubulin (Ac tub, gray). (F and H) MCCs stained for ODF2 (magenta), VANGL2 (magenta) and centriolin (cyan) (F), acetylated tubulin (gray), VANGL2 (gray) (H) or ODF2 (magenta). Scheme indicates the location of the basal bodies, acetylated microtubule crescent and/or VANGL2 and the angle between these structures. (G and I) Circular plots depicting the respective orientation of VANGL2 and basal feet (G), VANGL2 and acetylated microtubule crescent (I). n, number of analyzed cells: (G) n=94; (I) n=55. (J) Quantification of MCCs with aligned or misaligned basal bodies (BB) in cells with or without acetylated microtubule (Ac tub) crescent. n, number of analyzed cells: n=168.

**Supplemental Figure S3, related to Figure 3.**
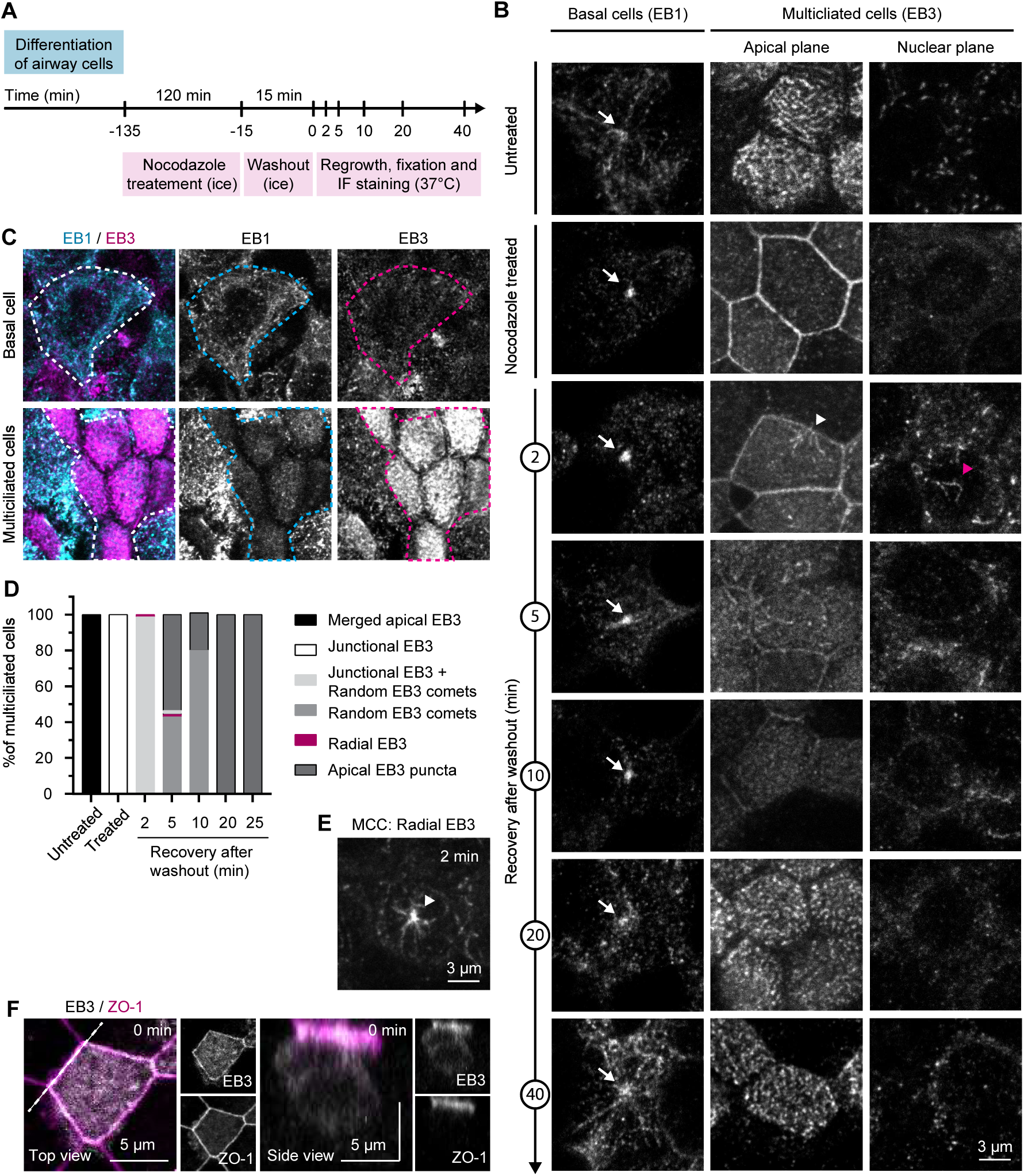
Microtubule nucleate from basal bodies in nocodazole washout assays. (A) Scheme of nocodazole washout assays. Abbreviation: immunofluorescence (IF) staining. (B) Microtubule regrowth after nocodazole washout in basal cells and MCCs stained for EB1 and EB3, respectively. Optical sections through the apical plane and the cell nucleus, representative for each microtubule recovery stage, are shown. White arrow shows centriole; white arrowhead shows microtubule nucleation from the cell-cell junction, magenta arrowhead indicates an EB3 comet. (C) Presence of EB1 (cyan) and EB3 (magenta) in basal cells compared to MCCs. Dashed line shows outline of a basal cell (top) or a patch of MCCs (bottom). (D) Quantification of MCCs with a particular EB3 localization pattern in nocodazole washout assays categorized into apical EB3, junctional EB3, junctional EB3 + random EB3 comets, random EB3 comets, radial EB3 and apical EB3 puncta. n, number of analyzed cells: untreated, n=17; treated, n=17; 2 min, n=71; 5 min, n=77; 10 min, n=43; 20 min, n=145; 25 min, n=106. (E) Examples of microtubule regrowth 2 min after nocodazole washout in MCCs stained for EB3. Arrowhead indicates microtubule nucleation in a radial pattern. (F) Top view (left) of an MCC fixed immediately after nocodazole washout and stained for EB3 (white) and ZO-1 (magenta). Dashed line represents the plane shown on the right.

**Supplemental Figure S4, related to Figure 3.**
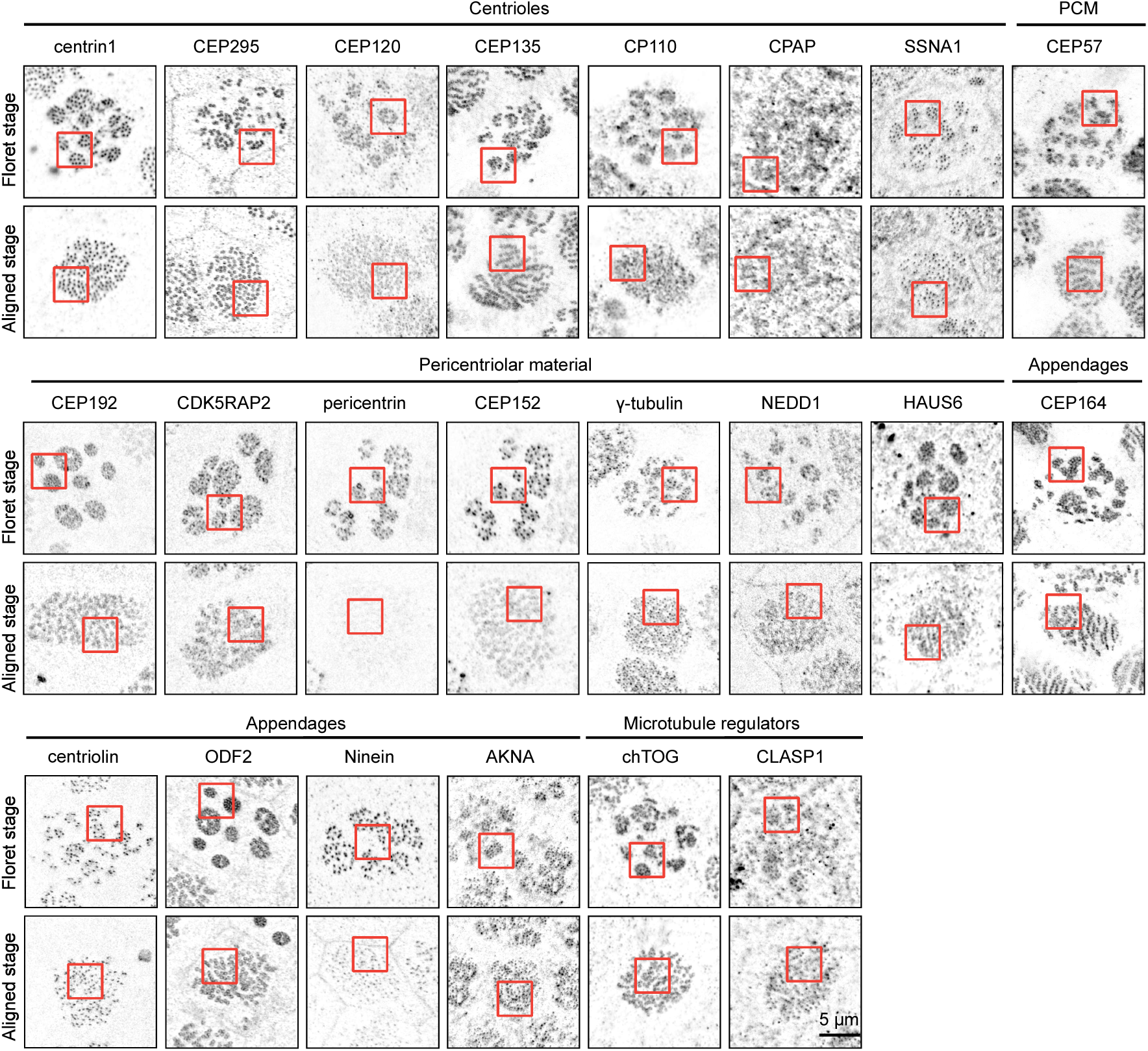
Imaging of basal body proteins using STED microscopy. Basal bodies in cells with a floret or aligned basal body pattern stained for various proteins, acquired using STED microscopy. Boxes indicate regions shown in Fig. 3E.

**Supplemental Figure S5, related to Figure 4.**
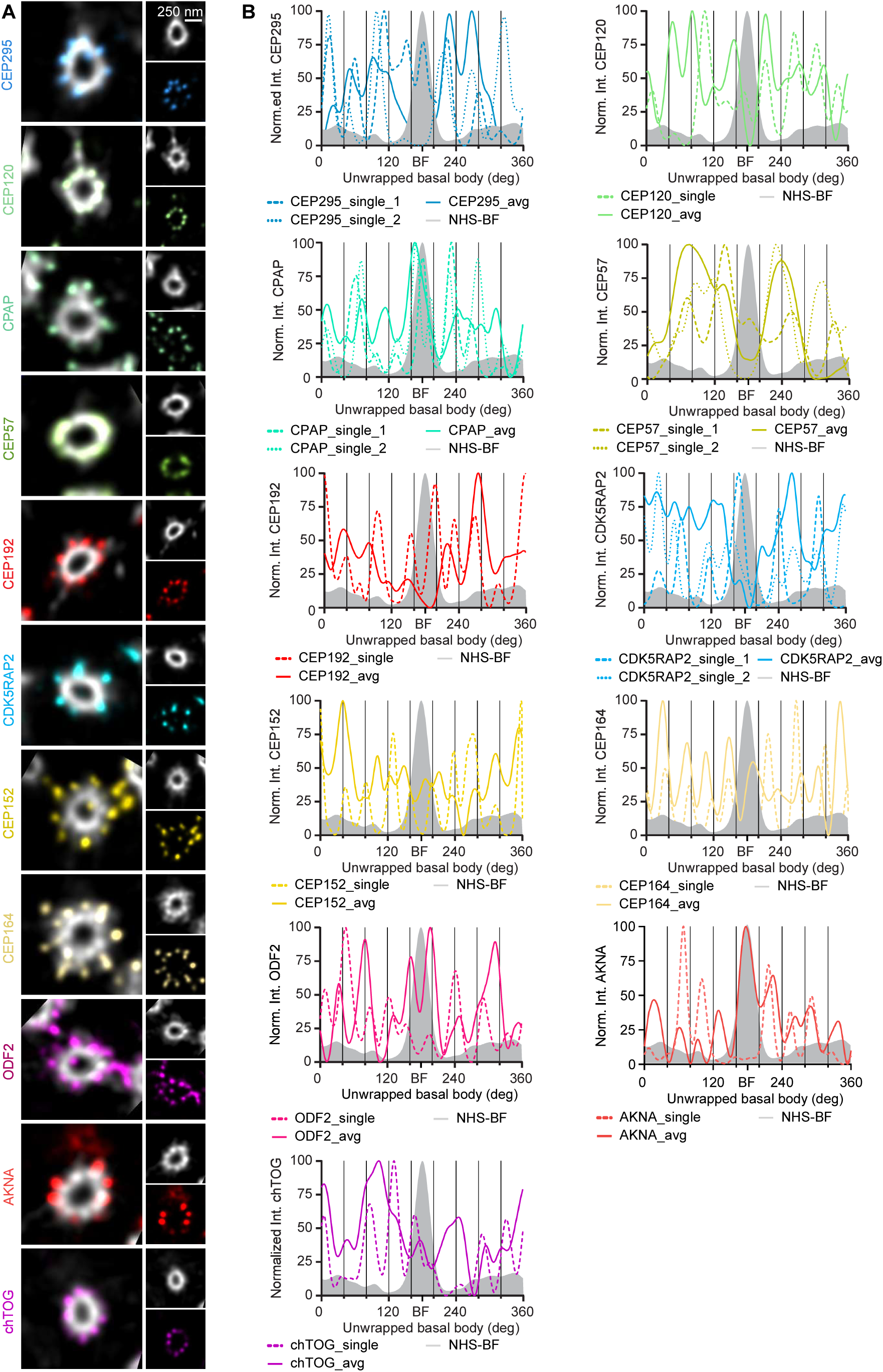
Polar transformation of ring-like organized basal body proteins. (A) Representative non-averaged, deconvolved images of basal bodies stained for various proteins (colors) and NHS ester (white) and imaged using TREx to show the symmetry of ring-like protein localization. (B) Angular intensity distribution of various proteins (color) and averaged basal body (NHS ester, gray). For proteins of interest, continuous line represents measurements performed on a basal body average, dashed line represents measurements performed on single basal body. X axis shows polar transformation from 0-360°, where vertical lines are placed at 40° increments characteristic for the nine-fold symmetry.

**Supplemental Figure S6, related to Figure 5.**
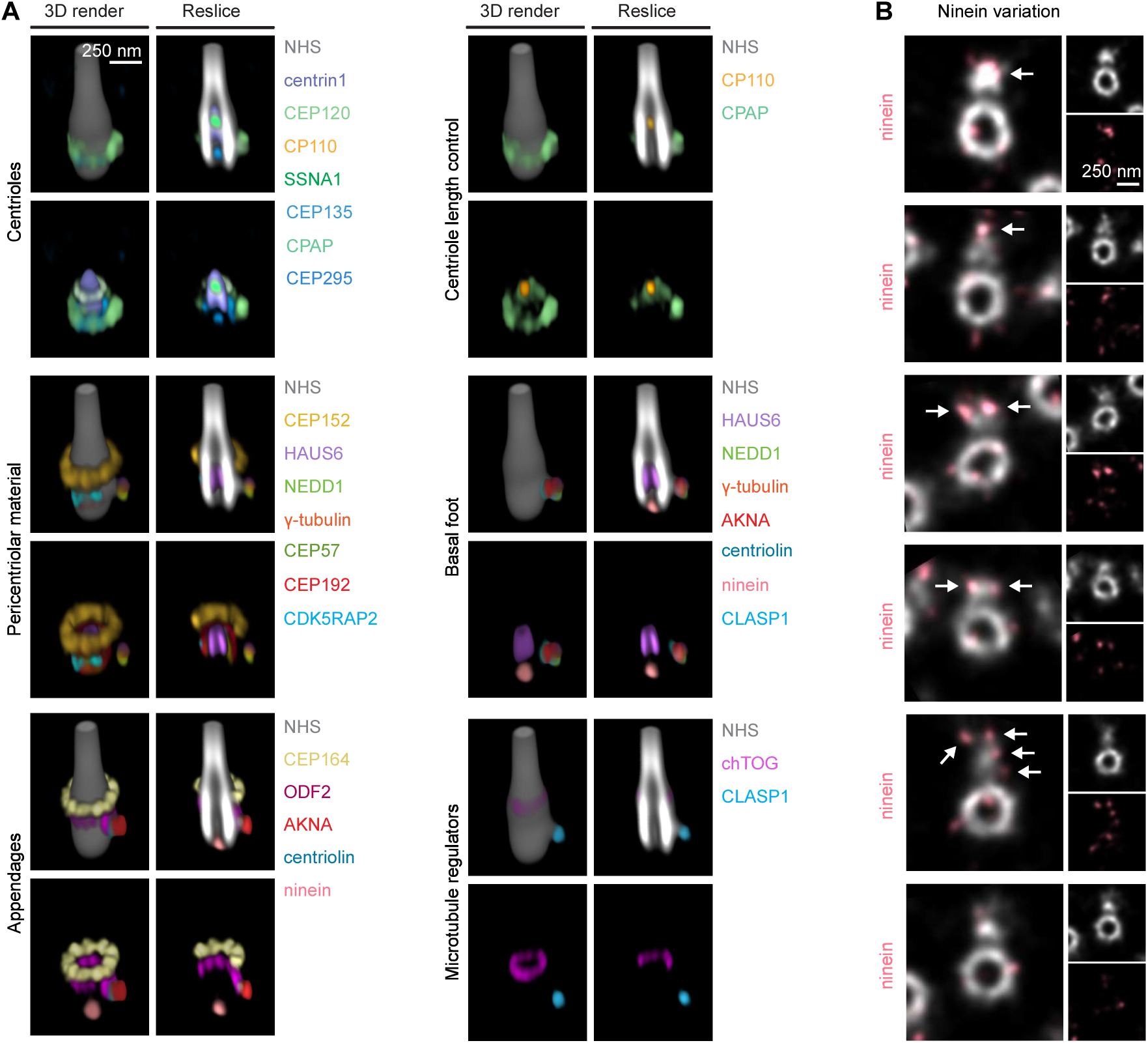
Combined architectural map of the basal body. (A) Gallery of basal body architectural maps of protein combination. The basal body average (NHS ester) is shown in white, and various proteins are displayed in colors. (B) Representative, non-averaged, deconvolved basal bodies stained for ninein (pink) and NHS ester (white) and imaged using TREx. White arrow points towards ninein localization on the basal foot.

**Supplemental Figure S7, related to Figure 7.**
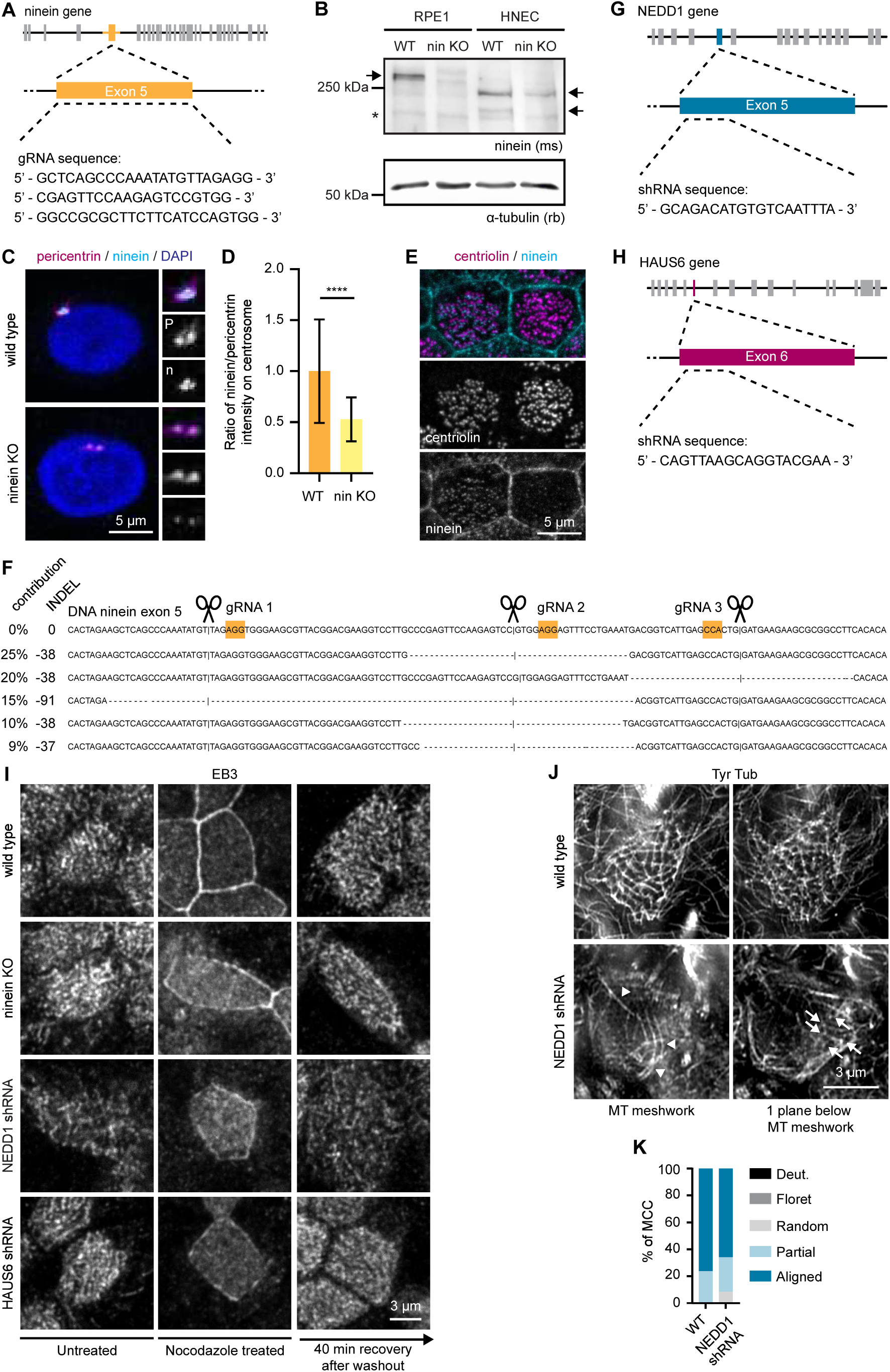
Generation and characterization of ninein knockout and NEDD1 and HAUS6 depleted basal cells. (A) Scheme of ninein gene with the location of three gRNAs used to generate ninein KO cells. (B) Western blot illustrating ninein KO in human nasal epithelial cells (HNEC) basal cells, and RPE1 cells as control. Arrows point towards ninein in RPE1 and HNECs; asterisk indicates a background band. (C) Basal cells of wild type and ninein KO cells stained for ninein (cyan), pericentrin (magenta) and DAPI. (D) Normalized ratio (mean±SD) of fluorescence intensity of ninein and pericentrin staining at centrosome in wild type and ninein KO basal cells. n, number of analyzed cells, from 3 independent experiments: wild type, n=335; ninein KO, n=639. ****P<0.001 was calculated using unpaired two-tailed Mann-Whitney U test. (E) Basal bodies of wild type and ninein KO cells stained for ninein (cyan) and centriolin (magenta) to show variation of ninein expression in ninein KO MCCs. (F) Sequencing results of the ninein KO cells using gel-purified PCR products. Wild type ninein exon 5 sequence (top) and genome edited ninein with different INDELs and their contribution in the cell population. Sequencing results of most common (70% of total) INDELs are shown. PAM sites of gRNAs (yellow) and cut sites (scissors) are indicated. (G,H) Schemes of NEDD1 and HAUS6 genes with the locations of shRNAs used for NEDD1 (G) or HAUS6 (H) depletion. (I) Microtubule regrowth after nocodazole washout in MCCs stained for EB3. Comparison between wild type, ninein KO, NEDD1 shRNA and HAUS6 shRNA-expressing cells. (J) Z-plane with apical microtubule (MT) meshwork and z-plane 0.3 µm below microtubule meshwork of MCCs stained for tubulin (white) and imaged using Airyscan2. In NEDD1 shRNA cell, arrowheads show microtubules associated with cell-cell junctions, and arrow indicates microtubules associated with basal feet. Comparison between wild type and NEDD1 shRNA cells. (K) Quantification of MCCs in wild type and NEDD1 shRNA condition categorized into deuterosome (Deut.), Floret, Random, Partially aligned (Partial) and Aligned basal body pattern. n, number of analyzed cells: wild type, n=42; NEDD1 depleted cells, n=59.

## Supplemental Movies

**Movie 1, related to Figure 1G. 3D render of a basal cell stained for tyrosinated tubulin.**

**Movie 2, related to Figure 1H. 3D render of a secretory cell stained for tyrosinated tubulin.**

**Movie 3, related to Figure 1K. 3D render of an MCC ALI day 5 stained for α-tubulin.**

**Movie 4, related to Figure 1L. 3D render of an MCC ALI day 10 stained for α-tubulin.**

**Movie 5, related to Figure 1M.: 3D render of an MCC ALI day 20 stained for α-tubulin.**

**Movie 6, related to Figure 2D. 3D render of an MCC ALI day 10 stained for tyrosinated tubulin (magenta) and acetylated tubulin (cyan).**

**Movie 7, related to Figure 2E. 3D render of an MCC ALI day 30 stained for tyrosinated tubulin (magenta) and acetylated tubulin (cyan).**

**Movie 8, related to Figure 5. Architectural map of the basal body.**

## METHODS

### CONTACT FOR REAGENT AND RESOURCE SHARING

Further information and requests for resources and reagents that are not related to human donor material should be directed to and will be fulfilled by the Lead Contact, Anna Akhmanova (a.akhmanova.uu.nl). Requests for use of patient samples and cell lines derived from them can be directed to Jeffrey M. Beekman.

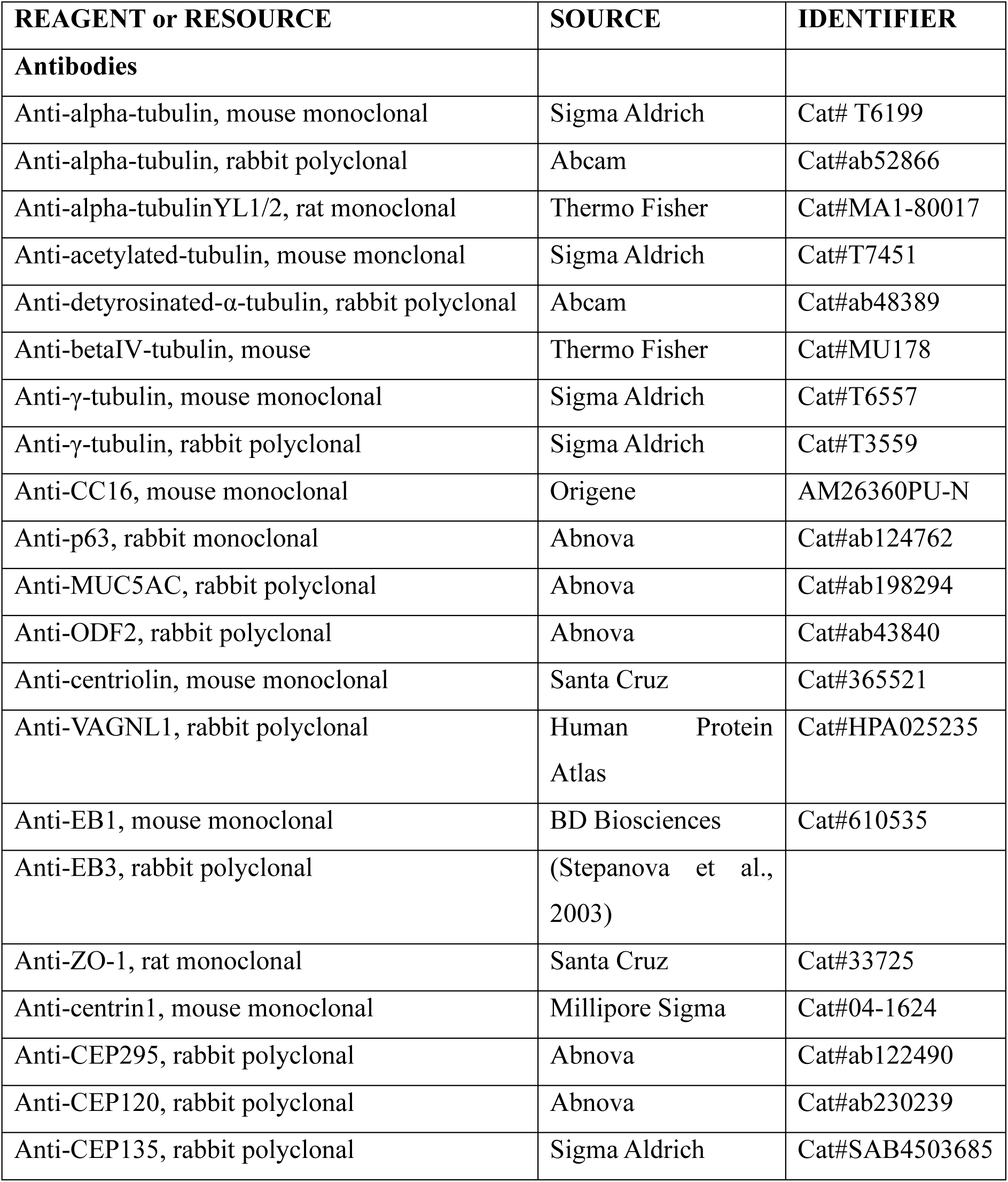
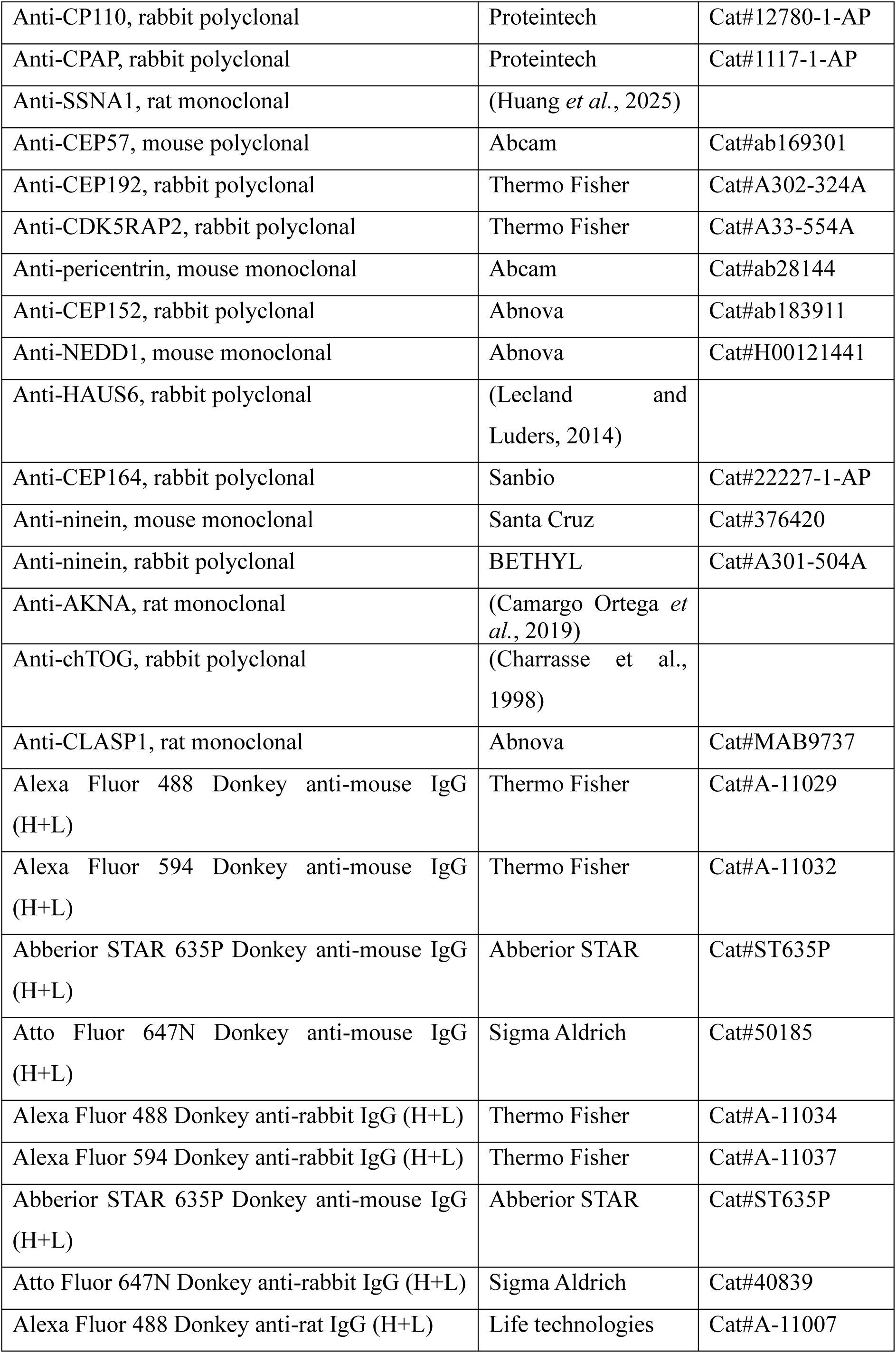
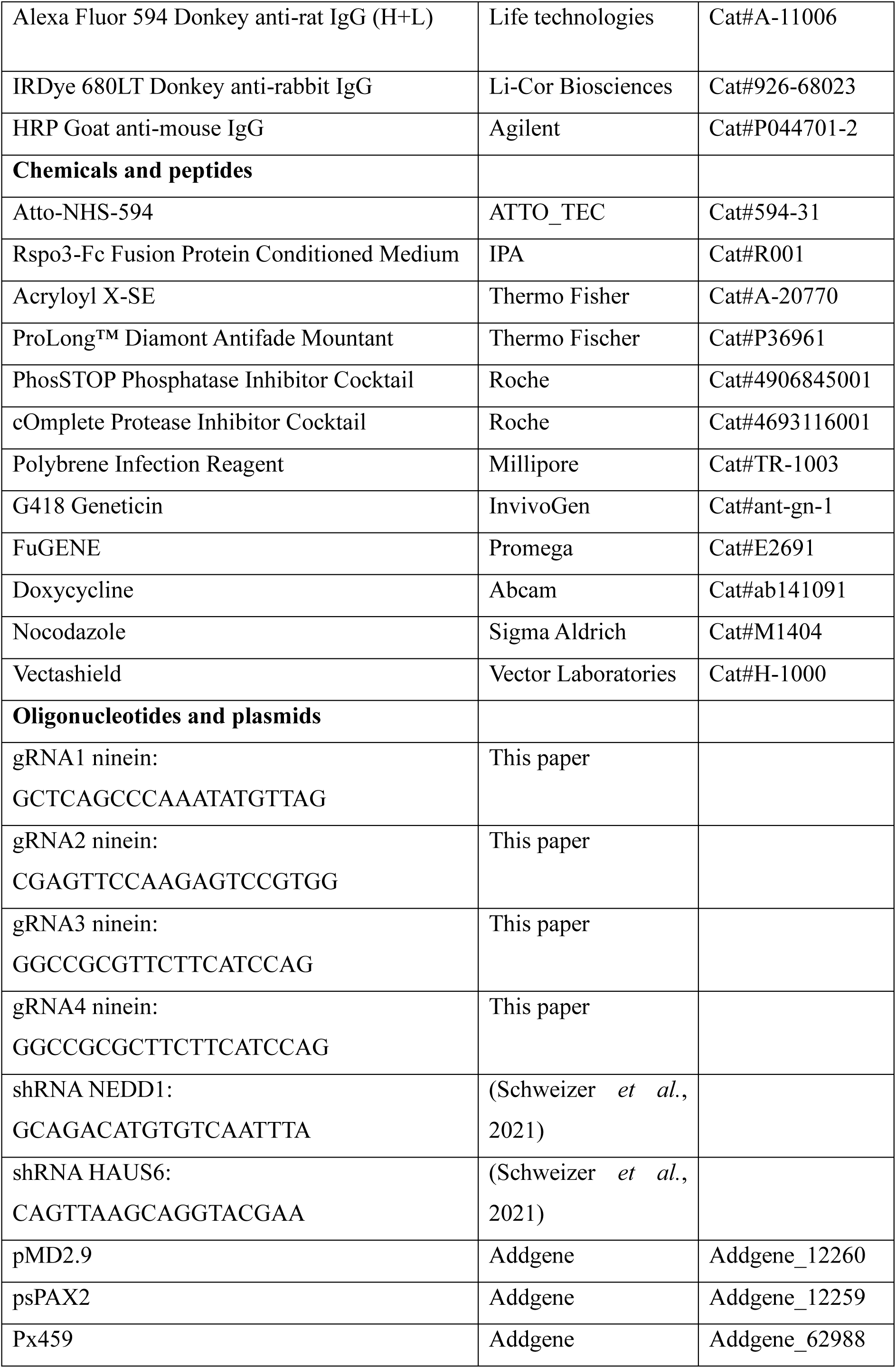
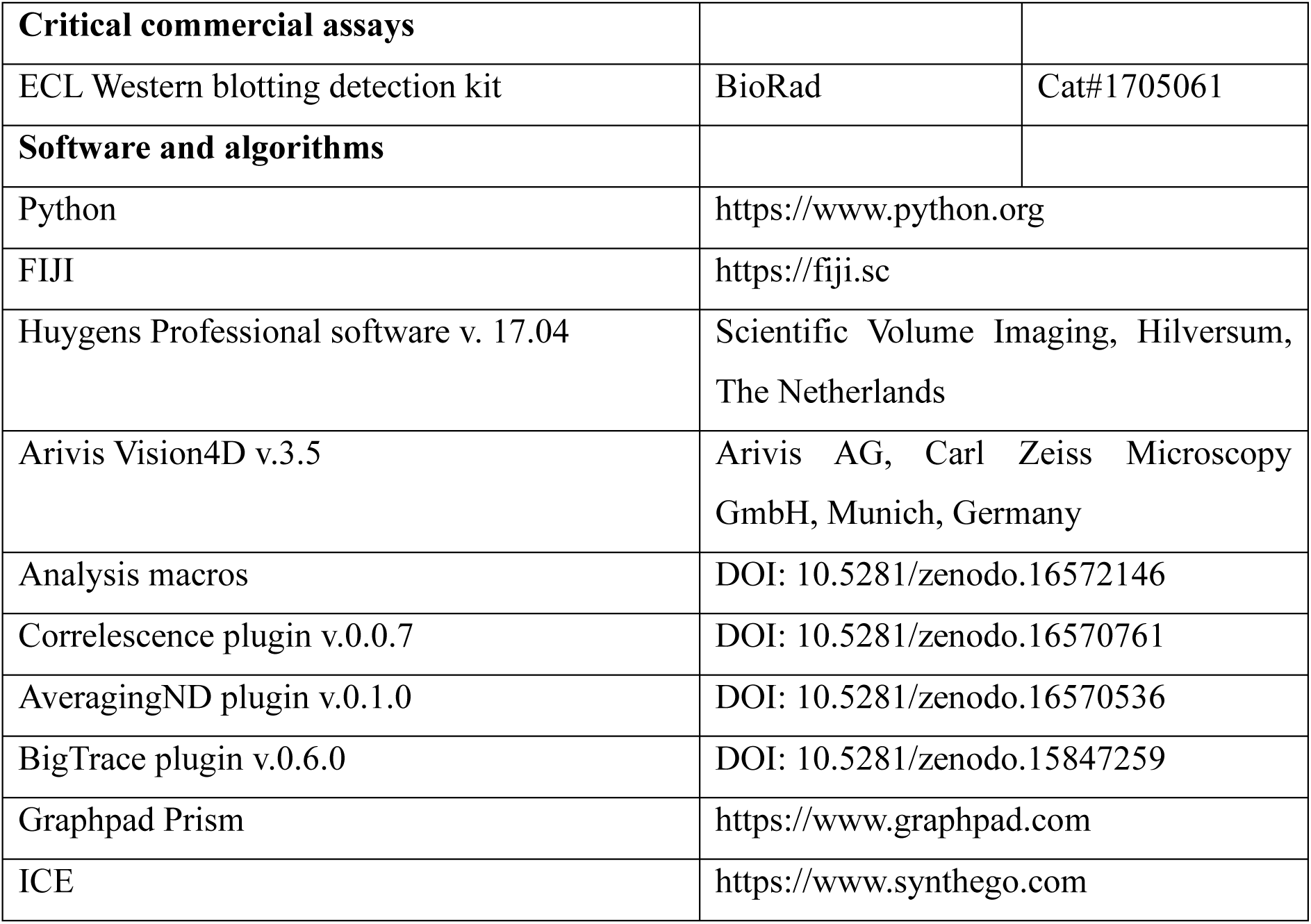

#### Cell culture RPE-1 cells

RPE-1 and RPE-1 ninein knock out cell lines were cultured in 50% DMEM and 50% F-10 supplemented with 10% fetal bovine serum and 1% penicillin/ streptomycin.

#### Cell culture of Human Nasal Epithelial Cells (HNECs)

Nasal brushes were collected from healthy human patients; patient samples gave explicit broad consent for biobanking in the UMC Utrecht Airbank (approved by ‘Toetsingscommissie biobanking (TcBio)’ protocol ID 16-586). Specific use for this study was approved under protocol (Tcbio ID 22-079). In this study, materials of two Human Nasal Epithelial Cells (HNEC) donors, HNEC4 and HNEC3, were used. The HNECs were collected and differentiated as described (Amatngalim et al., 2022). In brief, HNEC basal cells were expanded on 50 μg/ml collagen-IV-coated 6-well plates and cultured in basal cell (BC) expansion medium (50% BEpiCM-b, 23.5% Advanced (Ad) DMEM/F12, 20% Rspo3-Fc Fusion Protein Conditioned Medium, 10 mM HEPES, 1% GlutaMAX, 1% penicillin/streptomycin, 2% B-27 supplement, 0.5 μg/mL Hydrocortisone, 100 nM 3,3,5-triiodo-L-thyronine, 0.5 μg/mL Epinephrine Hydrochloride, 1.25 mM N-Acetyl-L-cysteine, 5 mM Nicotinamide, 1 μM A83-01, 1 μM DMH-1, 5 μM Y-27632, 500 nM SB202190, 25 ng/mL HGF, 5 ng/mL hEGF, 5 μg/mL DAPT, 25 ng/mL FGF-7 and 100 ng/mL FGF-10). For differentiation of 2D air-liquid interface (ALI)-HNEC cultures, basal cells were dissociated using TrypLE, seeded onto 30 µg/mL PureCol-coated 24-well transwells and cultured under submerged conditions in BC medium. Upon reaching confluency, the BC medium was changed to ALI-differentiation medium (50 nM A83-01, 0.5 ng/mL hEGF, 100 nM 3,3,5-Triiodo-L-Thyronine, 0.5 μg/mL Epinephrine hydrochloride, 100 nM TTNPB (Retinoic acid agonist), 0.5 μg/mL Hydrocortisone and 1% penicillin/streptomycin in 492,5 mL AdDMEM/F12) supplemented with additional 500 nM A-83. After 3-4 days under submerged conditions, cells were cultured in 42 μl ALI-differentiation medium supplemented with A-83 to minimally submerge the cells. After cells reached a compacted state, differentiation was triggered in ALI-differentiation medium supplemented with 5 µM DAPT under minimal submerged conditions for ∼15 days. Medium was refreshed 2 times a week; during differentiation the apical side of the cells was washed once a week using 125 μl phosphate buffered saline (PBS).

Unless otherwise indicated, experiments were performed on HNECs cultured on ALI for at least 20 days.

#### Lentivirus production and generation of stable transgenic cell lines

Lentiviruses were produced by a MaxPEI-based co-transfection of envelope vector pMD2.9 and packaging vector psPAX2 (Addgene_12260 and Addgene_12259). In brief, the supernatant of the packaging cells was harvested up to 72 h after MaxPEI-based transfection, filtered through a 0.45-μm filter and incubated with a polyethyle glycol (PEG) 6000-based virus precipitation buffer at 4°C overnight. Virus suspension was then centrifuged at 1500x g for 30 min at 4°C, the supernatant was removed, and the lentivirus was resuspended in PBS.

HNEC03i basal cells (van Grinsven et al, in preparation) were plated onto 12-well plates to reach 60% confluency at day of infection. Before infection, cells were incubated with 5 μg/mL Polybrene Infection Reagent for 1 h. Cells were infected twice with 2.5 μl lentivirus. To select cells with viral integration, HNEC infected with lentivirus were grown in presence of 500 μg/mL Geneticin (G418).

#### Western Blotting

For Western blotting, RPE-1 cells and basal cells were lysed in RIPA buffer (50 mM Tris HCl pH 7.4, 150 mM NaCl, 1% Triton X-100, 0.5% Sodium deoxycholate, 0.1% SDS, 1 mM EDTA and 10 mM NaF) supplemented with 1x protease inhibitor cocktail and 1x phosphatase inhibitor. Samples were incubated at 98°C for 10 min before loading onto an SDS-PAGE gel and transferred onto a nitrocellulose membrane using wet transfer for 2 h at 400 mA. Then, the membrane was blocked in 3% bovine serum albumin (BSA) for at least 1 h and incubated in primary antibody diluted in 3% BSA overnight at 4°C. Antibodies were removed by 3 washes of 10 min in TBST (0.05% Tween-20 in Tris-Buffered Saline (20mM Tris, 150 mM NaCl in MiliQ (TBS)). To visualize the loading control, α-tubulin, the membrane was incubated with IRDye680CW donkey anti-rabbit antibody in 3% BSA for 1 h at room temperature, washed in TBST as described before and exposed on the Odyssey CLx infrared imager (Li-Cor Biosciences). To visualize ninein, that has a weak signal that required a sensitive protein detection method, the membrane was incubated with horseradish peroxidase (HRP)-labelled secondary antibody in 3% BSA for 1 h at room temperature and washed in TBST as described before. Enhanced chemiluminescence (ECL) substrate was added to detect chemiluminescent and membranes were images on Image Quant LAS 4000.

#### Generation of CRISPR/Cas9 knockout cell lines

Generation of the ninein knockout basal cells was performed by electroporation of HNEC3i with Cas9/sgRNA mixture. In brief, 2.5 μl Cas9 (20 μM), 8.3 μl multi-guide sgRNA (a mix of gRNA1, 2 and 3 indicated at 30 μM in Reagent or Resource table) against ninein and 14.2 μl OPTIMEM supplemented with Y-27632 (10 μM) were premixed on ice. 0.5 million basal cells resuspended in a mix containing 1% B-27 supplement, 0.5 mM glutamine, 15.6 μM glutamate and 1% penicillin/streptomycin in Neurobasal medium were added to the protein/sgRNA mixture to make a total volume of 100 μl. Then, cells were electroporated 2 times with a 20 ms 1500 Volt pulse using an Invitrogen™ Neon™ Transfection System and 100 μl tips. Immediately after electroporation, cells were plated in a 12-well plate and cultured for 24 h in BC expansion medium supplemented with 10 μM Y-27632 and in the absence of penicillin/streptomycin. The RPE-1 ninein knockout cell line was generated by transfection of px459-sgRNA4 using FuGENE 6.

#### Doxycycline treatment

To induce shRNA expression, airway cells were cultured in the presence of 5 μg/mL doxycycline for a total of 30 days: 15 days of differentiation with DAPT, and a subsequent 15 days while air-exposed.

#### Nocodazole treatment

Prior to nocodazole treatment, HNECs were washed with PBS for 5 min, the filters were carefully cut from the transwell chambers and sectioned into pieces using a scalpel. The filter pieces were incubated in a 24-well plate with 0.5 mL ALI differentiation medium supplemented with 20 μM nocodazole with constant agitation for 2 h on ice. Nocodazole supplemented medium was refreshed after 1 h to ensure complete disassembly of the microtubule network. Nocodazole washout was carried out by 8 quick wash steps with 0.5 mL ice-cold wash medium (10 mM HEPES, 1% GlutaMAX, 1% penicillin/streptomycin, in AdDMEM/F12) on ice. Then, 0.5 mL 37°C pre-warmed ALI differentiation medium was added to the filter pieces and incubated in a 37°C incubator. After regrowth for the indicated time, cells destined for immunofluorescence staining with antibodies against EB proteins were fixed using ice-cold MeOH for 20 min at –20°C followed by post-fixation in 4% paraformaldehyde and 4% sucrose in PBS for 10 min at room temperature. Cells processed for immunofluorescence staining with antibodies against tubulin and subsequent Airyscan2 imaging were fixed in 4% paraformaldehyde and 4% sucrose in PBS for 20 min at 37°C. For the ‘nocodazole treated’ condition, cells were fixed immediately after the ice-cold washout. After fixation, the cells were processed for immunofluorescence as described below. When nocodazole washout experiments were performed on cells grown in presence of doxycycline, the nocodazole treatment and microtubule regrowth was performed in presence of 5 μg/mL doxycycline.

#### Immunofluorescence cell staining

Prior to fixation, HNECs were washed with PBS for 5 min. When one filter was used for multiple experiments, the filter was carefully cut from the transwell and sectioned into pieces using a scalpel; all fixation and wash steps were performed in a 24-well plate. For immunofluorescence staining with antibodies against centriolar proteins, cells were fixed in ice-cold MeOH for 20 min at –20°C. In case antibodies against EB proteins were used, MeOH fixation was followed by fixation with 4% paraformaldehyde and 4% sucrose in PBS. Cells destined for immunofluorescence staining with antibodies against tubulin were pre-extracted using 0.2% glutaraldehyde and 0.35% Triton X-100 in MRB80 (80 mM PIPES pH 6.8, 1 mM EGTA and 4 mM MgCl2) for 2 min at 37°C, followed by post-fixation in 4% paraformaldehyde and 4% sucrose in MRB80 for 20 min at 37°C. In all other cases, cells were fixed in in 4% paraformaldehyde and 4% sucrose in PBS for 20 min at 37°C. Fixation was followed by 3 PBS washes of at least 10 min each. In case of fixation with paraformaldehyde alone, samples were permeabilized using 0.5% Triton X-100 for 30 min or 1 h in case a staining against tubulin was performed. Then, cells were blocked in 3% BSA in PBS for at least 1 h at room temperature. Filter pieces were incubated in a 0.5 mL Eppendorf tube with 30 μl primary antibody dissolved in 3% BSA at 4°C overnight with constant agitation. Then, cells were washed by 3 PBS washes of 10 min and incubated in secondary antibody as described for primary antibody incubation for at least 3 h at room temperature. After 3 PBS washes of 10 min, cells were dehydrated in 70% EtOH followed by 100% EtOH and mounted in ProLong Diamond or Vectashield using 13 mm thickness 1.5 coverslips.

#### Ten-Fold Robust Expansion (TREx)

Ten-Fold Robust Expansion (TREx) microscopy was performed as described by (Damstra *et al*., 2022). In brief, cells were fixed and stained as described above. After removal of the secondary antibody solution, cells were post-fixed in 4% paraformaldehyde in PBS for 10 min at room temperature and incubated in 0.1 mg/mL acryloyl X-SE and 0.01% Triton X-100 in PBS overnight at room temperature with constant agitation. If samples were stained against tubulin, double concentration acryloyl X-SE was used. After 2 PBS washes of 15 min, filter pieces were placed onto a parafilm-covered glass slide with cells facing up, covered by a plastic transwell scaffold and sealed with grease. To this gelation chamber, 75 μl gelation solution (1% w/v sodium acrylate, 14.4% w/v acrylamide, 0.009% N,N’-methylenebisacrylamide, 1x PBS, 0.1% Triton X-100, 0.15% tetramethylethylenediamine (TEMED) and ammonium persulfate (APS) in MiliQ) was added and allowed to polymerize for 1 h at 37°C. The gel was carefully removed from the transwell scaffold and the gel section containing the filter piece was trimmed using a scalpel. Then, the gel was transferred to a 12-well plate, washed 2 times 15 min in PBS followed by homogenization in digestion solution (0.5% Triton X-100, 0.8 M guanidine-HCl, 9 U/mL Proteinase K in TAE (40 mM Tris, 20 mM acetic acid, 1 mM EDTA in MiliQ)) for 4 h at 37°C. After homogenization, the gel was thoroughly washed 3 times 15 min with PBS and in case a total protein stain was used, transferred to a 24-well plate and incubated in 20 μg/mL ATTO-NHS-594 or ATTO-NHS-647N in PBS for 1.5 h at room temperature. During the last 15 min of incubation, 5 μg/mL DAPI was added. To expand the sample, gels were transferred to a 15 cm petri dish with MiliQ and incubated at room temperature. After 30 min, the MiliQ was refreshed, and the sample was expanded overnight. Prior to imaging, the cells were trimmed and mounted onto a plasma-cleaned, poly-L-lysine treated cover glass.

#### Image acquisition and data processing

##### Confocal imaging

Data shown in Fig. 1C, 1I, 3C, 7A, 7B, 7E, 7F, 7I, 7J, S2A-C, S2E, S2H, S3B, S3C, S3E-G, S7C, S7E, S7I were imaged using a Leica TCS SP8 STED 3X microscope with continuous 405 nm and pulsed (80 MHz) white-light lasers, PMT and HyD detectors and spectroscopic detection with a HC PL APO 86x/1.20W motCORR STED (Leica 15506333) water objective.

##### STED imaging

Data shown in Fig. 1E, 3E, S2D, S2F and S4 were acquired using a Leica SP8 STED 3X microscope with 405 nm and pulsed (80 MHz) white-light lasers, PMT and HyD detectors and spectroscopic detection using a HC PL APO 93x/1.3 GLYC motCORR STED (Leica 11506417) glycerol objective. For STED imaging of Alexa Fluor 594, Atto674N and Abberior STAR635p, we used 594 nm and 633 nm white-light laser lines for excitation and a 775 nm synchronized pulsed laser for depletion. For multicolor STED imaging each fluorescent channel was imaged using the 2D STED configuration in line sequential z-stack, starting at the longer wavelength and ending at the shorter wavelength to prevent photobleaching.

##### Expanded sample imaging

Expanded samples shown in Fig. 1G, 1H, 1K-M, 2A-E, 3A, 3B, 3D, 4B, 4C, 4G, 5, S1A, S1B, S1D, S1F, S5A, S6A and S6B were imaged using a Leica TCS SP8 STED 3X microscope with continuous 405 nm and pulsed (80 MHz) white-light lasers, PMT and HyD detectors and spectroscopic detection with a HC PL APO 86x/1.20W motCORR STED (Leica 15506333) water objective.

##### Airyscan2 imaging

Data shown in Fig. 7M and S7I was imaged using a Zeiss LSM980 Airyscan 2 microscope equipped with 405 nm, 445 nm 488 nm 561 nm and 639 nm lasers, Axiocam 305 mono R2, 5.07 megapixel, 2464×2056, SONY IMX264 CMOS sensor with a Alpha Plan-APO 100x/1.46 OIL objective.

#### Quantifications of microtubule organization

##### Microtubule density maps

To estimate the number of microtubules along the longitudinal cell axis and generate the microtubule density map (Fig. 1N), a running maximum intensity Z projection of two slices was made of the expanded cell. Then, a maximum intensity projection of 2 Z-slices, followed by a Gaussian blur (1.0) was made at 5 selected positions along the longitudinal cell axis where 100% represent the apical side and 0% the basal side. The intersecting microtubules at each slice were counted and grouped into vertical (<0.15 μm), oblique (0.15-0.4 μm) and horizontal microtubules (>0.4 μm) (Fig. S1D,E).

##### Tracing of microtubule network with BigTrace

The FIJI (Schindelin et al., 2012) plugin BigTrace (v.0.6.0) was used to semi-automatically trace microtubules in images acquired using expansion microscopy. The traced lines 3D ROIs were used to extract quantitative information on the number, length, position and orientation of microtubules within the network (Fig. S1F,G). Signal intensities of microtubule subsets were measured on traced microtubules which were straightened using BigTrace. In the histogram of the position of the apical and basal microtubule ends, the uppermost 0.34 μm region at the apical side of the cell, where the microtubule meshwork is located, was excluded (Fig. 2G,H).

##### Fluorescence intensities of basal body components

For analysis of protein levels during MCC maturation, cell images acquired by STED microscopy were categorized into floret, random or (partially) aligned stages. A Gaussian blur (1.0) and background subtraction (Rolling ball, radius 50.0) were applied to a XY maximum intensity projection of 3 slices containing the basal bodies of one cell. Then, auto-thresholding was performed, and the selection was converted to a mask. The mask was used to measure the mean gray value on the unprocessed MAX projection. Intensity values of each protein of interest were normalized to the average intensity of the same protein at the floret stage.

##### Quantification of fluorescence intensities in genome edited or shRNA expressing cells

FIJI (Schindelin *et al*., 2012) was used for quantification of fluorescence intensities; all images of one experiment were acquired using the same microscope settings. A sum intensity projection was made comprising all z-slices with basal bodies. The region of interest (ROI) containing the basal bodies within one cell was drawn manually and was used the measure integrated intensity. Fluorescence intensities of γ-tubulin, NEDD1 and HAUS6 were normalized to the area comprising the basal bodies. In addition, due to high background signal with ninein antibody, ninein intensity was corrected for background: the background integrated intensity of ninein measured in a region without basal bodies was subtracted from the ninein integrated intensity measured in each MCC (corrected for the size of the MCC).

##### Apical microtubule meshwork

To quantify apical microtubule meshworks, images acquired with Airyscan2 were processed and binarized. A maximum intensity projection was made comprising the apical microtubule network and the microtubule filaments were enhanced using FIJI’s Tubeness plugin (Sato et al., 1998) with σ=5.7. The Tubeness-enhanced images were further thresholded using FIJI’s build-in auto-threshold tool with Otsu’s method and obtained binarized images underwent skeletonization with FIJI’s Skeletonize3D plugin. The integrated intensity within one cell was measured, and signal intensities were normalized to the cell area.

#### Generation of a 3D map of averaged protein densities at the basal body

##### Averaging of basal bodies per cell

The processing workflow to generate a normalized 3D map of averaged protein densities at the basal body was implemented as a set of FIJI macros and plugins. The raw and deconvoluted microscopy data, together with intermediate analysis images and plots are available at DOI: 10.24416/UU01-R1MHVT. The source code for the averaging analysis (including detailed description illustrated in Fig. 4A) is available at DOI: 10.5281/zenodo.16572146.

The input dataset consisted of confocal volumetric images (z-stacks) of apical surfaces of fixed and expanded HNECs. Each image contained a “total protein stain” NHS channel and one or two additional channels with the protein-of-interest staining. The XY size of the volume was set to include a surface of one cell, usually in the range of 50 x 50 μm. The range along z-axis was selected to fully include the basal body, transition zone and the beginning of the axoneme, in the range of 5-15 μm. For all images, the XY pixel size was equal to 46 nm and the z-step of 160 nm. The z-stacks were subjected to a mild deconvolution using Huygens Professional software version 17.04 (Scientific Volume Imaging, The Netherlands) with classic maximum likelihood estimation (CMLE) algorithm with parameters of Signal-to-Noise Ratio (SNR) equal to 10 over a maximum of 10 iterations.

Usually, we observed around 50-150 basal bodies per one cell surface. They were easily recognizable in the “total protein stain” channel as continuous tubular structures positioned along the apicobasal z-axis. The stained structure consisted of the basal body itself as a hollow cylinder of a fixed width with a short basal foot protruding in a perpendicular direction from the outside surface. It was followed by its gradually narrowing region, named the transition zone. Further in the apical direction, the transition zone becomes an axoneme, a long thin filament of a constant width. Individual structures were slightly bent away from the z-axis in a random direction, since they were caught by fixation at the moment of beating. To build an averaged volume, these structures needed to be transformed to a common similar homogeneous shape. The following paragraphs describe the procedures we developed for this “shape registration”. Briefly, the relative positions of the basal body and the transition zone enable us to identify the unique orientation along the z-axis. The protruding basal foot defines the rotation angle in the XY plane. Finally, if the bending deformation is removed, the transformed intensity shapes can be registered to each other without any assumption on the underlying symmetry.

All analysis and measurements described below were performed using the “total protein stain” channel as a reference. The channels with “protein-of-interest” staining were subjected to the same transformations as the reference channel.

At the beginning, we set out to determine an approximate XY position of individual basal bodies at an arbitrary Z-plane. Cross-sections of the cylindrical shape of a basal body in a single Z-plane looked like circles of approximately equal diameter. We used a crop of a single basal body circle cross-section as a template and utilized the FIJI multi-template matching plugin (Thomas and Gehrig, 2020) to detect position of multiple basal bodies in a user-selected single Z-plane. As an output, we got multiple rectangular 2D ROIs centered at basal body locations.

Inspection of the volumetric data showed that the structure of interest was not straight, but in most cases slightly bent away from the distal-apical z-axis. Therefore, in the next step we determined the 3D coordinates of a centerline of the basal body structure, i.e., the line that passes through the central rotational axis of symmetry in the middle. For that, we generated a template Z-stack with the same XY pixel size, where each plane contained an image a circle from a set of concentric circles ranging from an estimated maximum diameter *d_max_* = 2.1 μm to a minimum diameter *d_min_* with a step of 0.1 μm. We aimed to simulate basal body cross-sections of different diameters. After drawing each circle with a line of one pixel width, all images were convolved with a 2D Gaussian filter. Gaussian filter’s standard deviation was equal to the 2D standard deviation of the microscope *SD_PSF_* = 180 nm. Taking this convolution into account, the minimum diameter value *d_min_* was assigned the value of 3\**SD_PSF_*, since the circles smaller than that would be indistinguishable from each other.

At the next step, we cycled through all the ROIs at basal body locations, and for each of them performed the following procedure. At the initial z-plane we cropped a rectangular area with the same center as ROI and calculated normalized cross-correlation (NCC) in space between the cropped region and all slices of the Z-stack template using Correlescence plugin v.0.0.7 [https://doi.org/10.5281/zenodo.4534715]. It generates a Z-stack with NCC as pixel values and XY coordinates corresponding to the shift between the crop and template images. From this NCC z-stack we found a voxel with a maximum NCC value. Its z-coordinate corresponds to the circle with the best fitted diameter, and the XY coordinate can be used to determine the center of the basal body cross-section. Then the macro would move one slice up (or down) and crop the next area with the center at the newly determined XY coordinate. The procedure is repeated for all Z-planes and stops if the displacement in the center coordinates between adjacent slices exceeds 2\**SD_PSF_*. Then the macro works on the next basal body ROI location. As a result, for each single initial basal body ROI we obtain a table with XYZ coordinates of its centerline.

The centerline coordinates were used to construct 3D “linetrace” ROIs with diameter of 4.8 μm in the format, readable by the BigTrace plugin v.0.6.0 [DOI 10.5281/zenodo.15847258]. This plugin allows “straightening” of pipe-like 3D ROIs of an arbitrary thickness. The straightening was performed in the following manner. First, a continuous spline representation of the centerline was constructed. Each series of X, Y and Z coordinates were smoothed out with a moving window of 5 points. The series were reduced to include only every 5^th^ point and the end points. The obtained reduced smoothed series were interpolated by a cubic spline curve with an assumption of the second derivatives being zero at the end points. The continuous smooth spline interpolation allowed us to resample the centerline along its curve length with an equidistant step equal to the minimum of the voxel size (X,Y size of 46 nm). In addition, it provided the tangent vectors of the centerline curve at each location. To straighten the pipe-like volume around, we mapped (translated and rotated) these tangent vectors to be distributed along and aligned with a single (Z) axis in a new “straightened” space. Each tangent vector defines unique normal (perpendicular) plane in original volume. Sampling intensity in these planes allows to continuously map the intensity to the new XY plane of the “straightened” space. The only problem is that the tangent vector alone is not enough to define a unique XY frame orientation, since frame’s XY basis vectors can be freely rotated around the tangent at each point, remaining perpendicular. To build a set of consistent continuous normal frames at the new sampling points, BigTrace plugin was used to perform a computation of rotation minimizing frames (Wang et al., 2008). This method allows building of a moving coordinate frame along a space curve (centerline) where the frame rotates around the tangent vector as little as possible from one point to the next. A new XY pixel grid with the same pixel size was constructed in each normal plane, and the intensity values at grid locations were calculated using trilinear interpolation from the raw volumetric data. As a result, we generated a set of “straightened” basal body filament volumes with the isotropic voxel shape.

The final missing step in the shape normalization was the rotation of basal bodies in XY plane around the main symmetry z-axis, so that all their basal feet were aligned and pointed in the same direction. For that, we first built a XY maximum intensity projection image. Then we applied to it 2D tubeness filter (Sato *et al*., 1998) with a standard deviation equal to *SD_PSF_* to enhance the appearance of the basal foot and basal body ring cross-section and to reduce the background. We determined the coordinates of the center and the diameter of the ring using the same procedure with the z-stack template and NCC as described above. Then we created a new Z-stack template that at each z-slice contained radial line segment, rotating with one degree step around the center. The segment starting point was placed outside the ring boundary and it was spanning outward for the length of a half of the ring diameter. This template simulated all possible basal foot circular positions. To determine the best match, we calculated the maximum of NCC between line segment template images and the tubeness enhanced maximum projection image, excluding values corresponding to the spatial shift. This gave us the angular location of the basal feet. At this point, we upscaled all volumes two times in XY planes (to improve the averaging later) and rotated them along the central axis in a way that all detected basal feet were pointing upwards along the Y-axis.

Once the shape homogenization stage was finished, we used AveragingND plugin v.0.1.0 [DOI: 10.5281/zenodo.16570536] to perform iterative averaging/fusion and registration of multiple basal body volumes from the same cell. This was performed as follows: as the first step, we created an initial average template by aligning all volumes at their centers and calculating an average intensity for each voxel. The smaller volumes were padded with zero values to fit a minimal bounding box including all volumes. During the first iteration, we registered each volume to this template by finding a maximum of zero-masked NCC (Padfield, 2012). Before the NCC calculation, intensity of the analyzed image was subtracted from the average template, to minimize effect of its own presence. This kind of registration accounts only for the translation in 3D with a single voxel precision. During the second iteration, we built a new template, using registered volume positions and repeated registration step. Additionally, at each step we recorded an average of maximum NCC values for each volume as a quality measure. The procedure was repeated over user-provided number of iterations and the results of the iteration with the best of NCC average were chosen as final. The plugin also reconstructs and saves intermediate templates and the final registered input volumes. We defined the maximum number of iterations as being equal to four, since larger values did not improve average maximum NCC.

At the final stage, we determined diameter and *X_center_*, *Y_center_* centerline coordinates at each Z-slice of each averaged volume using the same NCC circle detection procedure.

##### Shape normalization of average basal body images

Visual inspection of averaged per cell volumes showed variability in the shapes of basal bodies, specifically in the diameter, length (height) of basal body and transition zone. It can be attributed to the variability of the expansion factor among gels, local deformations inside gels and cell-to-cell variability. To enable relative comparison of proteins-of-interest spatial distributions acquired on different samples, we performed “shape normalization” procedure described below.

At the first step, we segmented the z-axis location of the basal body and the transition zone. For that, we used our observation that the averaged image of basal body cylinder had an almost constant intensity which dropped at the point where the transition zone starts. To find segments marking basal body locations, we made a maximum intensity projection image in YZ plane and calculated a gradient magnitude image using FIJI FeatureJ plugin function Edges (with a gradient scale of 3 pixels). An intensity increase at the top of the basal body, a decrease at the beginning of the transition zone and an increase at its end manifested themselves on the gradient image as bright horizontal edges. To detect their positions, we built intensity profile along a central vertical line (at *Y_center_* position) and detected corresponding maxima locations *Z_1_*, *Z_2_* and *Z_3_*. In case when there were more than three maxima, their positions on the image were inspected manually and only those corresponding to the selected shape landmarks were preserved. From these values we calculated the height of the basal body *h_BB_* = *Z_2_* – *Z_1_* and transition zone *h_TR_* = *Z_3_* – *Z_2_*.

Next, we made two XZ and YZ single slice cross-section images at *X_center_*, *Y_center_* coordinates. The walls of basal body cylinder appeared on them as bright vertical lines. By detecting maxima at the horizontal line intensity profile at the Z position in the middle of the basal body *Z_1_* +0.5\**h_BB_*, we calculated diameters along X *d_X_* and Y axes *d_Y_*.

For the shape normalization, we calculated the average height of a basal body *<h_BB_>*, the average height of a transition zone *<h_TR_>*, and the average diameter of a basal body *<d_XY_>* (among all *d_X_* and *d_Y_*). We divided each volume into two parts along the z-axis at the position *Z_2,_* where the transition zone begins. Each of them was rescaled (compressed/stretched) using *h_BB_/<h_BB_>* factor for the bottom part, containing basal body and *h_TR_/<h_TR_>* for the top part (transition zone + axoneme). After that, each volume was rescaled along X and Y axes using factors *d_X_/<d_XY_>* and *d_Y_/<d_XY_>*.

At this point, the shapes of basal bodies lost their distinctiveness and became uniform, but rescaling changed the physical voxel sizes of the individual volumes. We decided to assign final volume dimensions by using a reference measurement of the basal body diameter of 250 nm, determined by EM (Li *et al*., 2012; Vladar *et al*., 2016).

The mutual registration of shape-normalized averaged volumes was performed using the AveragingND plugin as described above. As a final result, it generated registered versions of each volume, optimally translated to match the new final averaged template. These registered volumes were used for the subsequent comparative analysis and quantification of spatial distributions of the proteins of interest. The final average of averaged basal body shapes was used to calculate the XY coordinates of the central common reference axis.

##### Quantification of the radial distribution of averaged intensities

For radial symmetry analysis of basal body averages, measurements were performed on the maximum intensity projections of the ∼3 slices with maximum fluorescence intensity. For single basal bodies, a deconvolved basal body image was selected when the basal body fulfilled the following criteria: round, representative, no signal from neighboring basal bodies. The FIJI Polar Transformer plugin was used to convert the image to polar coordinates. Fluorescence intensity was then measured along the y-axis (representing the angle of the unwrapped basal body) using line width of 10 px or 30 px, for basal body averages and single basal bodies, respectively.

##### Quantification of full width at half maximum (FWHM) of basal body average

The intensity profiles along the center-to-periphery and longitudinal basal body axis were measured on basal body averages. The original X,Y,Z volumes were resampled to the cylindrical system of coordinates R,θ,Z with the FIJI Radial Profile Angle plugin. The axis of the cylinder runs along the original z-axis with its XY coordinate equal to the fitted XY position of the centerline of the averaged basal body. For quantifications, we used the maximum radius R_max_ of 110 pixels (∼550 nm). The angle θ was chosen to be equal to 0° degrees at the position of the basal foot. Using the plugin, we generated several intensity profiles I_prof_(R,Z) with different angle θ integration ranges, defined by the start and end angles [θ_start_, θ_end_]. The intensity value at I_prof_(R_i_,Z_i_) was equal to the mean intensity of the perimeter of a concentric circle of radius R_i_ at the z-slice Z_i_, where we take only the circular segment defined by angles θ_start_ and θ_end_. The first profile plot (total) contained a full radial scan of the basal body, where θ was in the range of [0°, 360°], the second profile plot contained only the basal foot region [-30°, 30°], and the third plot excluded basal foot [30°, 330°]. These profile plots were used to automatically determine the position of the maximum intensity peaks along the radial and longitudinal coordinates and additionally find the width at the half of maximum. To detect the luminal position of proteins that were also present on the outer basal body surface, the radius value including only the luminal region was chosen. Due to interference from other protein positions, to detect the basal foot position of CEP164 and ninein and the distal ring position of AKNA, the position of the maximum intensity peaks and corresponding half maximum was manually determined. Quantifications were performed on three basal body averages obtained from three individual cells. Then, the positions of the maxima and the half maxima were averaged and plotted using GraphPad Prism, where NHS-signal was used as a reference for the basal body and its proximal end (lower half maximum) was set to 0.

## Data and code availability

The raw and deconvoluted microscopy data, together with intermediate analysis images and plots are available at DOI: 10.24416/UU01-R1MHVT. The source code for the averaging analysis is available at DOI: 10.5281/zenodo.16572146.

